# Uncovering Microbial Biosynthetic Potential with Genomic Context-aware Protein Language Model

**DOI:** 10.1101/2025.04.29.651206

**Authors:** Zixin Kang, Haohong Zhang, Chaoqin Liang, Ronghua Yang, Ying Ye, Hong Bai, Yonghui Zhang, Kang Ning

## Abstract

Microbial secondary metabolites, synthesized by biosynthetic gene clusters (BGCs), offer vast potential for biotechnological applications. Among BGC profiling techniques, computational detection methods face challenges, including time-consuming alignment and reliance on predefined profiles. To address these, we present BGC-Finder, an end-to-end pipeline utilizing protein language models for BGC detection and annotation from microbial genomes and metagenomes. This approach achieves remarkable increase in profiling speed of up to 100-fold, and employs genomic context-aware modeling to facilitate interpretable genetic essentiality assessment and large-scale BGC clustering. BGC-Finder outperformed traditional methods, successfully detecting 9.49% more biosynthetic-core genes and 27.70% more cytochrome P450s in 742 experimentally-validated BGCs. Notably, it retrieved 31 remote biosynthetic homologs from 210 polar marine metagenomes and identified 4,585 BGCs with 6,388 core genes from 256 fungal genomes. These findings highlight BGC-Finder’s capability to illuminate “microbial biosynthesis dark matter” (sequence-unrelated, function-similar biosynthetic enzymes) and expedite natural product discovery.

**Highlights:**

- BGC-Finder is an accurate and ultrafast pipeline leveraging protein language models (pLMs) to predict and annotate biosynthetic gene clusters (BGCs) from microbial genomes and metagenomes.
- The genomic context-aware model enables interpretable analysis: attention-driven identification of essential biosynthetic genes and embedding-guided BGC clustering.
- BGC-Finder sensitively retrieves remote homologous BGCs from both bacteria and fungi genomes, uncovering hidden ‘microbial biosynthesis dark matter’.
- We discovered a non-ribosomal peptide synthetase (NRPS) family, which involved into function-specific BGCs in two evolutionarily distant fungi.

## Introduction

In complex ecosystems, microbial communities produce diverse secondary metabolites to mediate microbe-microbe and microbe-host interactions [1, 2]. These metabolites are primarily synthesized by biosynthetic gene clusters (BGCs) - co-localized genomic regions encoding enzymes like polyketide synthases (PKS), non-ribosomal peptide synthetases (NRPS) and terpene synthase (TPS) [3]. Within BGCs, biosynthetic-core genes encode enzymes that catalyze foundational metabolic reactions, while accessory genes facilitate modification, regulation and transportation [4, 5]. Advances in computational genome mining, coupled with extensive microbiome sequencing data, have accelerated the discovery of uncharacterized BGCs across bacteria, fungi, archaea, and uncultivated taxa [6–13].

Despite these advantages, current BGC profiling methods face critical bottlenecks. Traditional methods like antiSMASH [14] define the most routine alignment-based approaches for detection, using HMMER [15] to compare sequences against established profiles. These underscore several challenges. To begin with, there is an imbalance between the exceptional growth of sequencing data and the time-consuming nature of HMMER. Profiling BGCs across thousands of metagenomes typically requires hundreds of CPU hours. Besides, due to the reliance on HMMER, methods require constructing separate profiles for different taxa (e.g., bacteria and fungi), limiting their broad applicability. A generalized model capable of profiling BGCs across diverse taxa could enable deeper insights into microbiome biosynthetic potential. Furthermore, the frequent mutation, genetic rearrangement and natural selection has driven extensive BGC divergences over long evolutionary periods, enhancing microbial biosynthetic diversity [16–19]. Since only a small fraction of BGCs is experimentally characterized, alignment-based approaches struggle to identify ‘microbial biosynthesis dark matter’, referred to sequence unrelated, yet function similar biosynthetic genes, based on one-dimensional (1D) sequence information. Last but not least, many biosynthetic genes even lack detectable Pfam domains (e.g., *NvfI, NvfL*, *cp1B* and *cp2B*) [20, 21], further complicating detection. Unlocking this unexplored diversity could accelerate drug discovery, such as for antibiotics, chemotherapeutics and immunosuppressants, necessitating more advanced methods that combine high sensitivity and computational efficiency.

Recent advance in large language models (LLMs) pretrained on biological data [22] offer promising for BGC profiling. Protein language models (pLMs) [23–27], trained to discern patterns in protein sequences, capture co-evolution information to establish sequence-function-structure relationships. For example, ESM [28], pretrained on millions of proteins, leverages its embeddings to enable protein structure prediction and remote homology detection [29]. So, pLMs can be facilitated to encode biosynthetic genes to help the model to incorporate three-dimensional (3D) structure information rather than 1D sequence information. However, for BGC detection problem *per se*, pLMs treat proteins as isolated entities, overlooking genomic contexts shaped by evolutionary processes (e.g., regulation, transport) [30]. Hwang et al. constructed gLM, a genomic language model to address this by modeling pLM embeddings using a contextual architecture [31]. Combining pLMs with gLMs could address current limitations by leveraging genomic-function interplay.

Here we introduce BGC-Finder, a context-aware method integrating pLMs and gLMs for BGC detection and annotation in microbial genomes and metagenomes. We evaluated its performance in BGC boundary detection, class prediction, and gene function annotation. BGC-Finder achieves state-of-the-art (SOTA) performance in all areas across bacterial and fungal genomes, processes data >100× faster than alignment-based tools, and also enables downstream applications including interpretable genetic essentiality assessment and large-scale BGC clustering. Applied to polar marine metagenomes and fungal genomes, it uncovered extensive “microbial biosynthesis dark matter” corroborated by sequence and structure alignment. Notably, it revealed a conserved NRPS family in two evolutionarily distant genera: *Fusarium* and *Aspergillus*, subsequently evolving into function-specific BGCs.

## Results

### Overview of BGC-Finder

BGC-Finder comprises four deep-learning models: two ESM2 models [23] of different sizes, a BGC-detector model and an BGC-annotator model (**Fig. 1b, c**). To incorporate genomic contexts, contigs are represented as structured sequences of gene embeddings generated by ESM2. The pipeline starts with open reading frame identification using Prodigal [32] for bacteria or AUGUSTUS [33] for fungi. Genes are then encoded by ESM2-6M and screened by the BGC-detector to map putative BGCs. These BGCs are then re-encoded using ESM2-650M. Gene embeddings are concatenated with product class embeddings, and further added with token-type embeddings, and gene position embeddings to enhance context modeling (see **Methods**) in contextual embedding layer. Then, a five-layer transformer encoder in the BGC-annotator transforms them into context-aware BGC embeddings, where attention mechanisms decode gene roles and aggregate information toward the product class embedding. The genomic contxual embeddings are fed into a fully connected layer to predict BGC product classes and gene functions, and their pairwise cosine distances are leveraged to measure BGCs’ similarity. Besides, gene-class attention scores from the transformer are used to assess genetic essentiality in BGCs **(Fig. 1b).**

**Figure 1.**
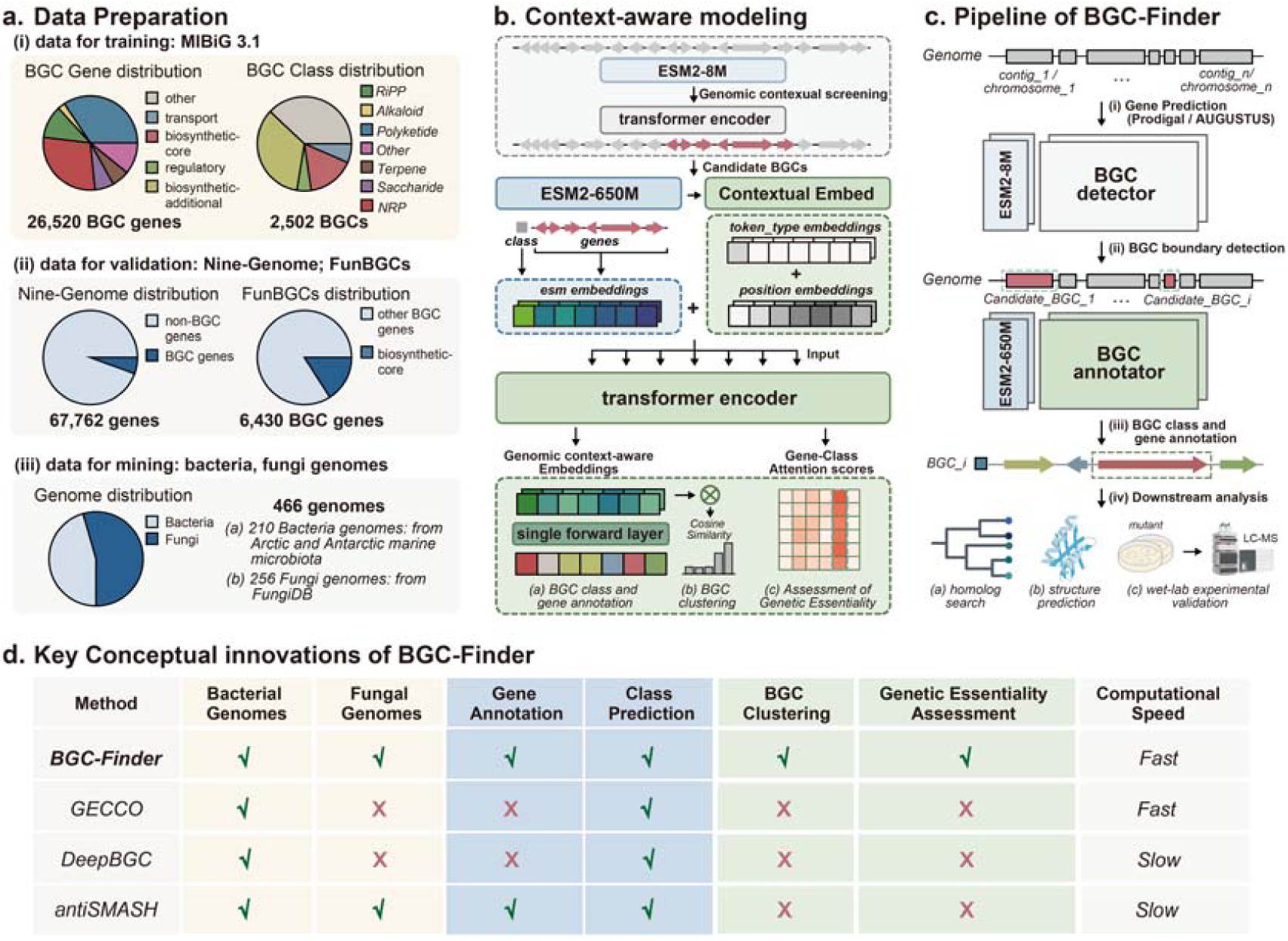
Overview of BGC-Finder. **a.** Data Preparation. (i) Data for model training was constructed using *MIBiG 3.1*, which contains 26,520 genes and 2,502 BGCs. (ii) Data for external validation comprised of two phylogenetically distinct datasets. A bacterial *Nine-Genome dataset* contains 67,762 genes with BGC-border annotation, and a fungal *FunBGCs* contains 6,430 BGC genes with function annotation. (iii) Data for mining also comprised 210 bacteria genomes from Arctic Antarctic marine microbiota and 256 fungal genomes from *fungiDB*. **b.** Context-aware modeling. The transformer encoder in BGC-detector screened BGCs from genomes based on genomic contexts. Candidate BGC genes were then re-encoded using ESM2-650M, with product class representation initialized as a zero vector. BGC-annotator was utilized to predict BGC class type and gene functions. Its embeddings enable similarity calculations, while gene-class attention scores quantify genetic essentiality within BGCs. **c.** Pipeline of BGC-Finder. Genes in bacterial genome were predicted using Prodigal, and genes in fungal genome were predicted using AUGUSTUS. BGC-detector was used to detect BGC boundary. Then, the BGC-annotator subsequently predicts product classes and gene functions, with prioritized candidate BGCs undergoing downstream analysis including homology searches, structure prediction, and experimental validation. **d.** Key conceptual innovations of BGC-Finder. The table summarizes the capabilities of various methods designed for BGC profiling. (Fast: 10^1^∼10^2^ second per genome; Slow: 10^2^∼10^3^ second per genome.)

To comprehensively capture the genetic patterns in BGCs, all experimentally validated BGCs from MIBiG 3.1 [34] were collected to train the BGC-annotator. This dataset included 26,520 genes and 2,502 BGCs sourced from bacteria and fungi **(Fig. 1a)**. The BGC-detector leverages our previous method, BGC-Prophet methodology [35]. Two phylogenetically distinct datasets from bacterial and fungal genomes were used to validate BGC-Finder’s performance in BGC boundary detection and gene annotation, while mining-sets were utilized to explore biosynthetic potential **(Fig. 1a)**. Novel BGCs was prioritized for further homolog searches, structure prediction and experimental validation **(Fig. 1c)**. Leveraging diverse datasets, we demonstrated that BGC-Finder can perform a range of profiling tasks across bacterial and fungal genomes, and achieve SOTA performance compared with current methods **(Fig. 1d)**.

### Generation of genomic contextual embeddings

Large-scale BGC clustering is vital for mapping microbial biosynthetic diversity and prioritizing BGCs. Conventional methods like BiG-SCAPE [36] and BiG-SLICE [37], rely on sequence alignment and predefined pHMM libraries to calculate BGC similarity. We validated that BGC-Finder’s genomic contextual embeddings enable alignment-free BGC representation and clustering.

To assess biological relevance, we first evaluated embedding regions were separated by BGC class. Embeddings from each BGC-annotator layer and ESM2’s last layer were extracted to determine the optimal layer. Represented as 1280-dimensional vectors, BGC pairwise similarity was measured via cosine distance. Deeper layers exhibited greater divergence in similarity distributions, with significant separation between same-class and different-class BGCs (P < 0.0001). UMAP visualization of final-layer embeddings showed that the BGC-annotator learned similar representations for BGCs with analogous biosynthetic mechanisms, while distinguishing metabolites within the same class. For instance, PKS and NRPS, which typically employ modular assembly-line mechanisms, clustered closely, whereas some non-typical BGC subclusters (e.g., NI-siderophore) diverged from main NRPS clusters. This indicates that, despite being trained with supervision, BGC-Finder learns an embeddings space reflecting both product class and specific biosynthetic mechanism without fine-tuning.

To further check whether embeddings’ similarity aligns with BGC similarity measured by other clustering approach, we compared embedding similarity to BiG-SCAPE scores. ESM2 embeddings showed the lowest correlation (Spearman’s ρ = 0.37), improving with layer depth (max ρ = 0.43). Notably, adding position and token-type embeddings to ESM2 outputs increased correlation (Δρ = +0.03), confirming enhanced context modeling.

### BGC-Finder outperforms alignment-based profiling approaches

Building on the validated capability for contextual embedding generation, we evaluated BGC-Finder against established methods using the Nine-Genome dataset [10, 12] for BGC boundary detection and the MIBiG test-set for gene function and class prediction. Firstly, to evaluate whether the model can effectively retrieve BGC genes while keeping a low false positive rate, precision and F_1_-score were utilized as index reflecting methods’ comprehensive performance (**Fig. 2b**). BGC-Finder outperforms all other methods in BGC boundary detection task. Then, we proved that BGC-Finder enables accurate BGC gene function annotation (AUROC = 0.990, AUPR = 0.986) in MIBiG test-set (**Fig. 2c**). For class prediction, it outperformed GECCO and DeepBGC in F_1_-score across all BGC class, excluding antiSMASH due to its role in MIBiG annotation (**Fig. 2d**). Besides, BGC-Finder also offers a significant advantage in computational efficiency. Alignment-based approaches typically require specialized hardware or software optimization and multiple independent runs, which requires 5 to 30 minutes to conduct genome-scale BGC profiling. BGC-Finder demonstrates processing speeds up to 10^2^-fold faster (**Fig. 2e)**.

**Figure 2.**
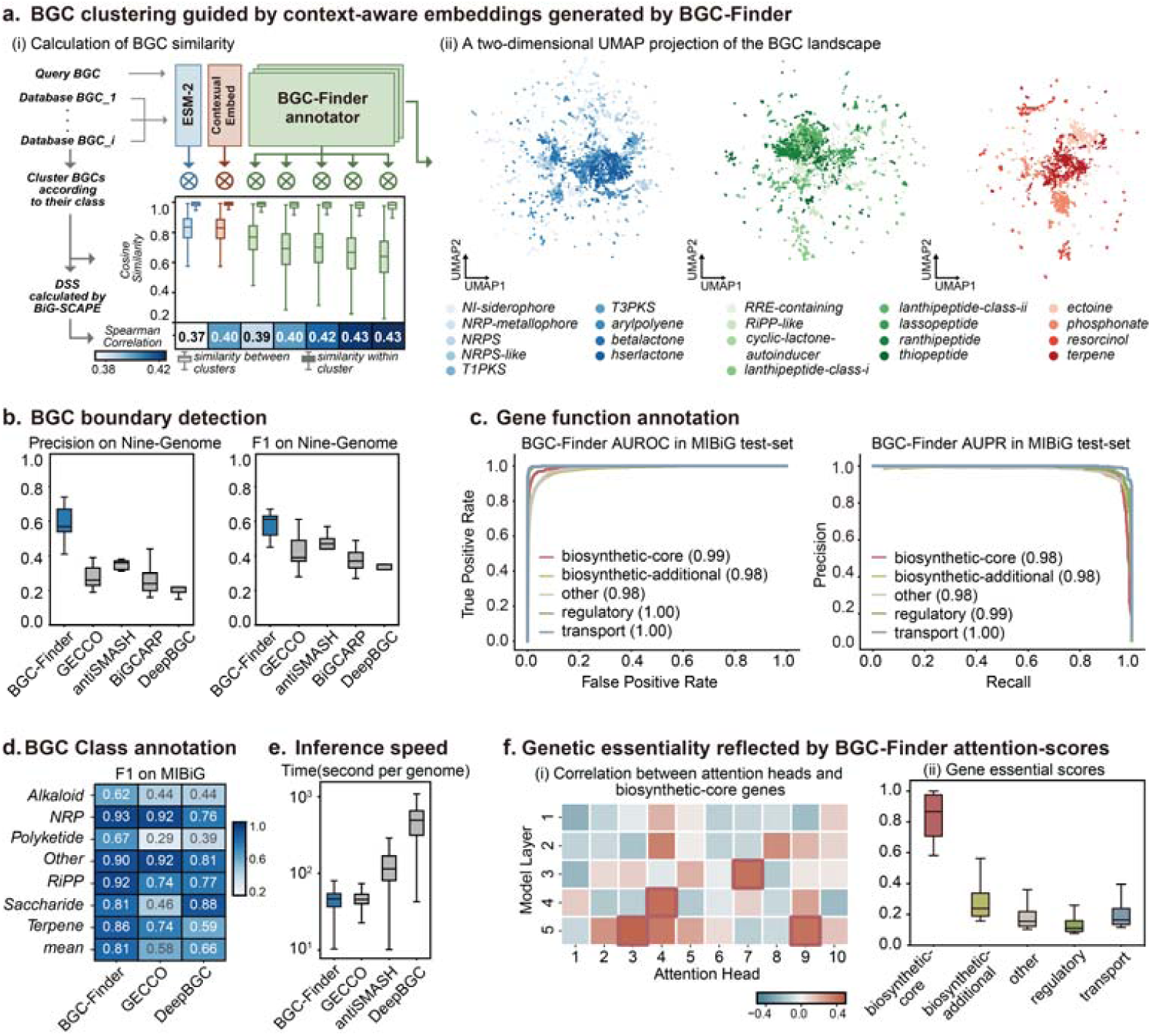
Performance evaluation and interpretable analysis of BGC-Finder. **a.** BGC clustering guided by context-aware embeddings generated by BGC-Finder. (i) The analysis involved generating embeddings for query BGCs and database BGCs using BGC-Finder’s BGC-annotator, followed by pairwise cosine distance calculations. Clusters were organized by product category, with intra-cluster and outlier similarities quantified. The strong Spearman correlation between model embeddings and BiG-SCAPE scores (DSS) validates the biological relevance of the embedding space. (ii) A Two-dimensional UMAP projection of the BGC landscape, representing 22,000 BGCs from main categories. **b-d.** Comprehensive benchmarking reveals BGC-Finder’s superior accuracy across multiple tasks. In BGC boundary detection (**b**), BGC-Finder achieves higher precision and F1-scores than GECCO, antiSMASH, BiGCARP, and DeepBGC when evaluated on the *Nine-Genome* dataset. For gene function prediction (**c**), the model demonstrates exceptional performance (AUROC = 0.990, AUPR = 0.986) on the MIBiG test-set. In BGC class prediction (**d**), BGC-Finder outperforms GECCO and DeepBGC in F1-score across all BGC classes. **e.** The computational speed of different methods. BGC-Finder is 2 orders of magnitude more computationally efficient than DeepBGC and antiSMASH. **f.** Attention mechanism analysis reveals biologically meaningful patterns in model decisions. (i) Spearman correlation between attention heads and biosynthetic-core genes. Red dot box marks attention head with Spearman correlation higher than 0.40. (ii) Gene essentiality scores derived from gene-class attention weights, where higher scores indicate a greater gene contribution to class inference. The top 5% of gene-class scores from each gene category were collected and displayed.

To assess rationality of our gene encoding strategy and model architecture, we further trained two Random Forest (RF) models: RF-Gene for gene function annotation task and RF-Class for product class prediction task. Performance comparison showed that the ESM2-embeddings-based RF model significantly outperformed its one-hot-encodings-based counterpart in both tasks (**Supplementary Fig. 3**). Furthermore, to validate biologically relevant distinctions captured by the embeddings, we observed that ESM2 embeddings exhibited a significant difference between core gene similarity and overall gene similarity (P < 0.0001), whereas no such difference was observed for one-hot embeddings (P = 0.51) (**Supplementary Fig. 4a, c**). Notably, while RF-Gene matched BGC-Finder’ s performance in gene function prediction, RF-Class’s average AUROC was only 0.791, even with ESM2 embeddings. This disparity underscores the advantage of BGC-Finder’s transformer-based architecture over traditional machine learning models in harnessing genomic features for class inference task.

Since BGC-Finder’s attention mechanism, we leveraged the attention scores learned by the model to explore whether it has captured genetic feature (**Fig. 2f)**. Several attention heads strongly correlated (Spearman’s ρ > 0.40) with core gene positions, indicating their roles in identifying essential gens. Furthermore, we collected all gene-class attention scores and categorized them by gene function to evaluate their contributions to class prediction (see **method**). Core genes showed the highest genetic essentiality, followed by tailoring (biosynthetic-additional) genes and transport genes, aligning with biosynthesis principle. Interestingly, transport genes showed a notable essentiality, consistent with established evidence [38]. These findings show BGC-Finder’s ability to extract key biosynthetic information from context embeddings and capture genetic essentiality without any prior biological knowledge.

### Precise identification of diverse biosynthetic genes

Expanding the scope of our evaluation, we applied predictive analytics to FunBGCs [20], a dataset containing 742 experimentally validated fungal BGCs with detailed function annotations. BGC-Finder outperformed antiSMASH in biosynthetic-core gene retrieval, achieving an accuracy of 0.96, precision of 0.95 (842/891), recall of 0.82 (842/1,025), and F1 score of 0.88 **(Fig. 3a, b)**. It accurately detected more genes across different categories, like geranylgeranyl pyrophosphate synthase and DMATS-type prenyltransferase (**Fig. 3b, Table S1**). Besides, many tailoring (biosynthetic-additional) genes misclassified by antiSMASH were correctly annotated by BGC-Finder. For instance, cytochrome P450 monooxygenase contributing to a broad range of biological functions in versatile metabolism [39, 40], were reported to frequently misclassified by antiSMASH [41], with 159 annotation errors out of 733 instances in FunBGCs. In contrast, BGC-Finder accurately identified 731 cases (**Table S1**). However, it failed to detect several enzymes responsible for new, non-canonical product synthase, like phosphoenolpyruvate mutase and isocyanide synthase. Besides, both BGC-Finder and antiSMASH identified terpene cyclase as tailoring genes rather than core genes.

**Figure 3.**
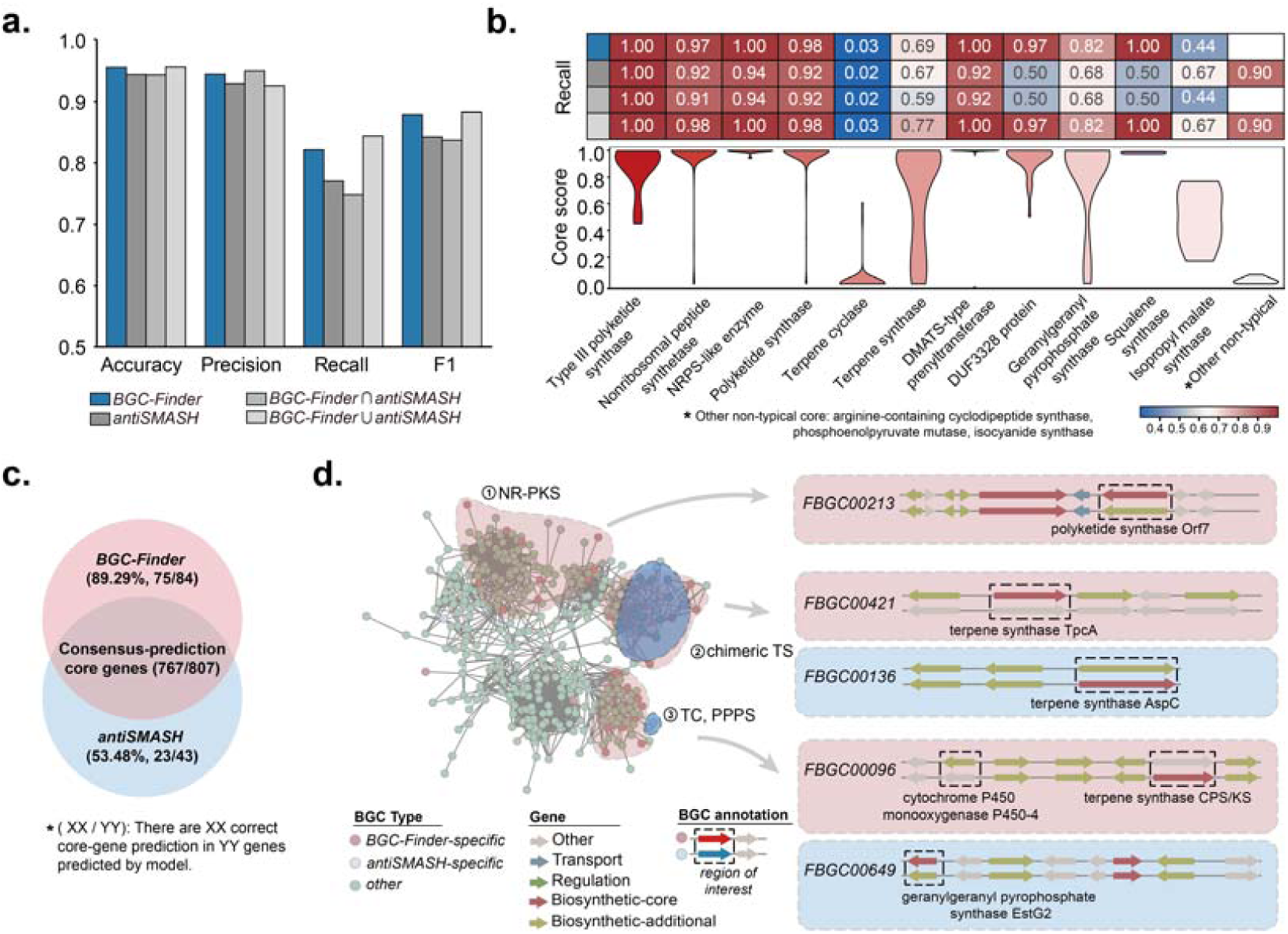
Comparative analysis of BGC-Finder and antiSMASH in detecting biosynthetic-core genes within experimentally validated fungal BGCs. **a.** Performance metrics of four strategy (BGC-Finder only, antiSMASH only, BGC-Finder n antiSMASH, BGC-Finder ∩ antiSMASH) in detecting core genes within experimentally validated fungal BGCs. **b.** Recall rates on different core genes and confidence score distribution of BGC-Finder for different core genes. **c.** Venn diagram of prediction overlap, with BGC-Finder detecting 842/1,025 validated core genes (Recall: 0.82) versus antiSMASH’s 790/1,025 (Recall: 0.77), and higher accuracy in uniquely identified genes (75/84 vs. 23/43). d. The BiG-SCAPE network ( dss > 0.35 ) highlighting BGCs with core genes exclusively identified by BGC-Finder (red) or antiSMASH (blue).

To further check BGC-Finder’s performance on core gene prediction, we examined cases where two models’ predictions overlapped and diverged. In the overlapped set of 807 core genes, 767 were validated correct **(Fig. 3c)**, yielding a higher precision of 0.95 compared to individual model predictions. The union of predictions (combining the overlapped and diverged sets) retrieved more core genes and achieved a higher recall of 0.84 (865/1,025) **(Fig. 3a**, **Fig. 3b)**. BGC-Finder’s superior performance is further exemplified by its ability to detect remote homolog missed by antiSMASH. For instance, in *FBGC00213*, antiSMASH misclassified a polyketide synthase as a tailoring gene, while BGC-Finder identified it as a core gene, corroborated by experimental validation. The core gene was validated as a remote homolog by querying it on Foldseek [42] server and aligning it with known enzyme sharing low sequence similarity (SI: 39.35%) but high structural similarity (TM-score: 0.51) (**Fig. 3d, Supplementary Fig. 5**). This highlights BGC-Finder’ s robustness in pinpointing key biosynthetic enzymes and its sensitivity in capturing remote homologs.

### Retrieval of remote biosynthetic homologs from polar microbiome

Having demonstrated BGC-Finder’s ability to accurately identify diverse biosynthetic genes, we applied the method to metagenomes from geologically isolated biomes: Arctic and Antarctic marine environments to further validate BGC-Finder’s ability to discover ‘microbial biosynthesis dark matter’ hidden in these unique niches by retrieving distantly homologous biosynthetic-core genes. 1,636 BGCs were identified and annotated by BGC-Finder and antiSMASH, of which 715 were both identified by two methods (overlap) and 894 were only identified by BGC-Finder (BGC-Finder-specific) (**Fig. 4a**). To verify whether BGC-Finder could capture more diverse core genes, we calculated sequence diversity across three groups: all core genes, core genes identified by BGC-Finder and core genes identified by antiSMASH (see **method**). The core gene sequence diversity of BGC-Finder was 0.04 higher than that of antiSMASH. Furthermore, we prioritized representative BGCs by clustering BGCs and selecting main clusters (n > 25) (**Fig. 4b**). In the two main clusters, BGC-Finder broadened five subclusters by retrieving 31 distantly core genes overlooked by antiSMASH. Foldseek was applied to search homologs of these core genes. The core genes in *subcluster_1,2,3* all involved in the terpene synthase. Specifically, based on its low sequence identity (SI: 20.9%) and extremely high structural similarity (TM-score: 0.845) with the homolog, we inferred that *k141_289389_12* from *Leeuwenhoekiella* is an isozyme of squalene-hopene cyclase, an enzyme involved in hopene skeleton formation. Besides, the core genes in subcluster_4,5 were involved in the fatty acid biosynthesis and shikimate pathway, respectively. The analysis revealed that BGC-Finder can discover hidden biosynthetic pathway by capturing shared structure information in biosynthetic remote homologs.

**Figure 4.**
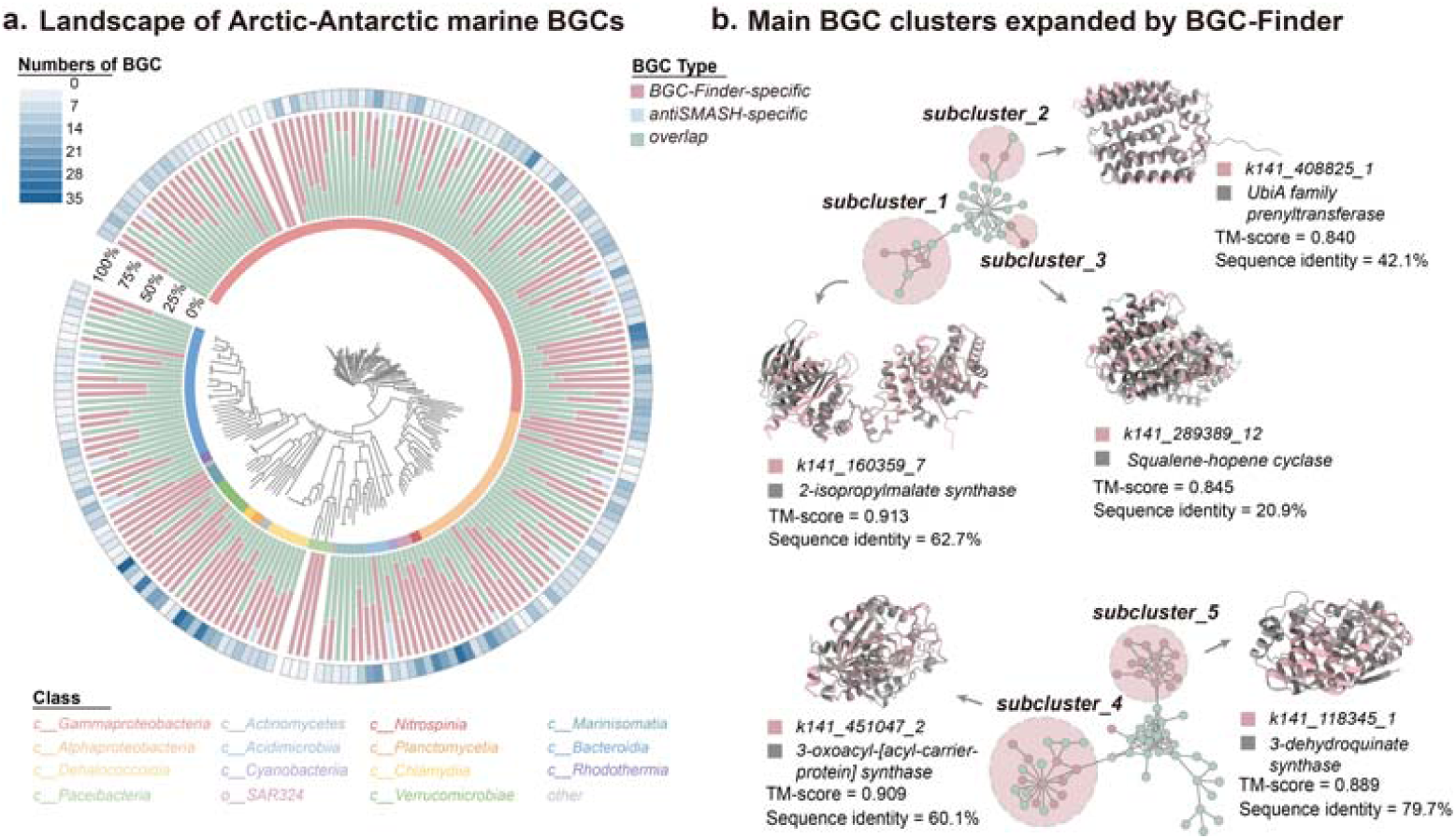
Sensitive retrieval of distantly biosynthetic-core genes and expansion of ‘microbial biosynthesis dark matter’ with BGC-Finder. **a.** Landscape of Arctic-Antarctic marine BGCs. BGC-Finder and antiSMASH were used to predict BGCs from Arctic-Antarctic marine microbiota MAGs. A phylogenetic tree was built for 210 bacterial MAGs based on a concatenated alignment of bacterial single-copy genes and placement of each MAG in the GTDB-Tk reference tree to present BGC distribution. The Inner rings depict BGC distribution: overlap (BGCs both identified by BGC-Finder and antiSMASH), BGC-Finder-specific (BGCs only identified by BGC-Finder) and antiSMASH-specific (BGCs only identified by antiSMASH). The outer ring illustrates the distribution of BGC counts across the 210 species. **b.** Main BGC clusters expanded by BGC-Finder. The network constructed by BGC similarity calculated by BiG-SCAPE. The main clusters (dss > 0.30, n > 25, dss: BGC similarity, n : number of BGCs) were extracted for further analysis. Here, BGC-Finder discovered distantly biosynthetic-core genes in four BGC subclusters (*subcluster_1, subcluster_2, subcluster_3, subcluster_4, subcluster_5*). Specifically, the gene k141_289389_12 from *Leeuwenhoekiella* was potentially isozyme of squalene-hopene cyclase.

### Comprehensive profiling of BGCs from fungal genomes

To showcase BGC-Finder’s ability to uncover hidden microbial biosynthetic potential, we analyzed 256 fungal genomes from 197 species across major phyla (e.g., *Ascomycota* and *Basidiomycota*) [43]. Leveraging the pipeline, we identified 6,388 core genes within 4,585 fungal BGCs (Fig. 5a).

**Figure 5.**
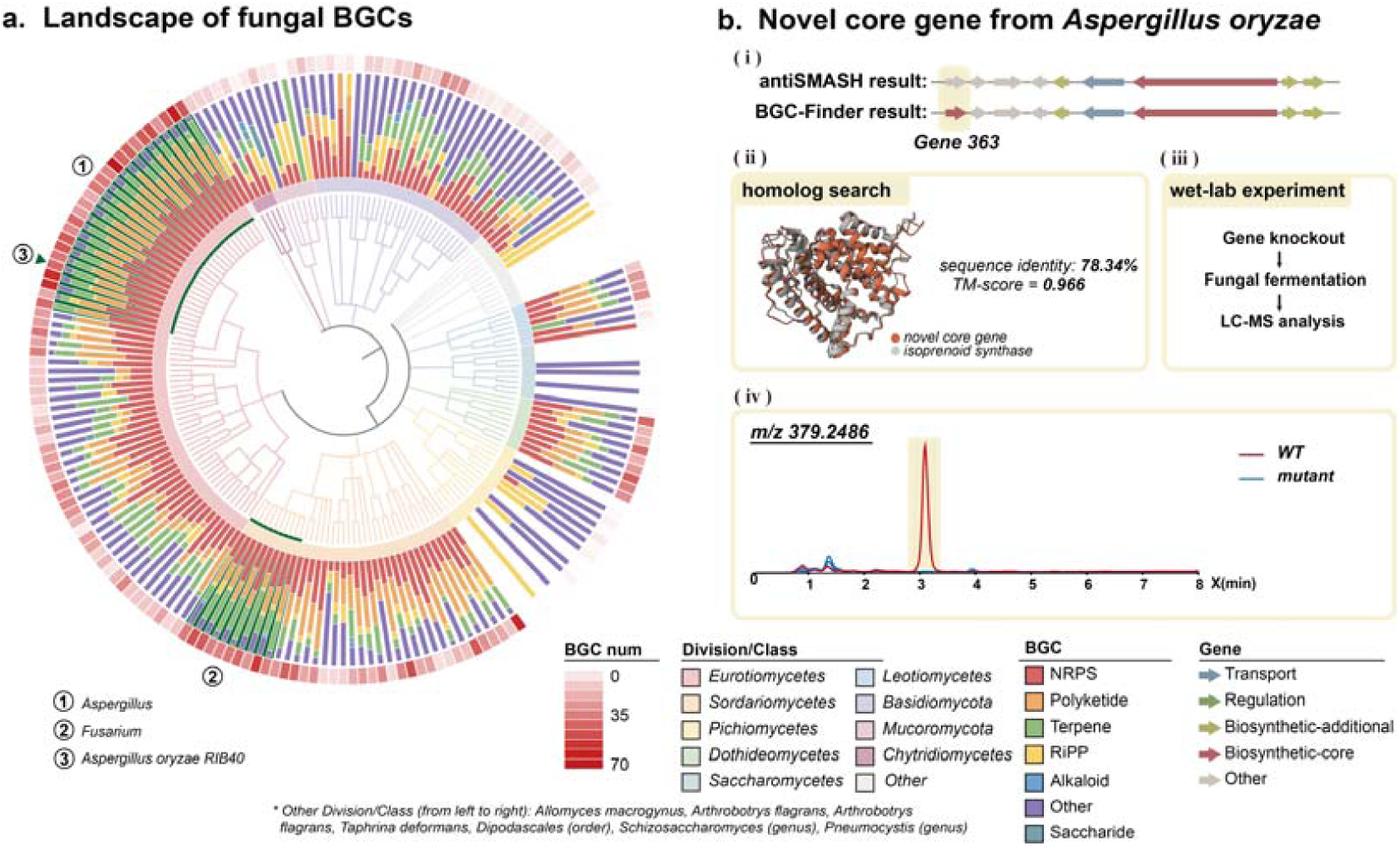
The global distribution of fungal BGCs and identification of a novel core gene. **a.** Phylogenomic tree constructed at the order level, encompassing 197 fungal species from the FungiDB database. Average numbers of biosynthetic gene clusters (BGCs) and core genes per genome are shown for each order. Inner rings depict BGC distribution across categories: NRP, polyketide, terpene, RiPP, saccharide, alkaloid, and others. The outer ring illustrates the distribution of BGC counts across the 197 species. **b.** (i) Comparison of BGC-Finder and antiSMASH annotations for an NRP BGC identified from Aspergillus oryzae. (ii) BGC-Finder identified an additional core gene, a terpene synthase, exhibiting high structural similarity (TM-score = 0.966) and moderate sequence identity (78.34%) to a known isoprenoid synthase. (iii) LC-MS analysis of metabolites extracted from rice cultures of WT and mutant A. oryzae strains lacking the novel core gene. Extracted ion chromatograms (EICs) at m/z 379.2486 reveal WT-specific peaks (highlighted in yellow), absent in the mutant, indicating products of the terpene synthase.

Fungi stand out among microbial producers due to their remarkable biosynthetic richness and ecological significance [44–46]. Despite this, the vast biosynthetic diversity of fungal BGCs remains underexplored, limiting potential application in biotechnology, agriculture, and medicine [3, 45, 47]. Beyond the experimentally validated BGCs cataloged in databases like MIBiG [34] and FunBGCs [48], many fungal BGCs lack functional characterization. This gap stems from the time-intensive and costly nature of wet-lab validation, compounded by the unique properties of fungal BGCs, such as remote homology and complex genomic contexts, all of which make them challenging to analyze [49].

BGC-Finder revealed a diverse array of BGCs, many of which were previously uncharacterized. Among the 6,388 core genes, 2,220 were annotated as ‘hypothetical’ or ‘unspecified’ by Gene Ontology annotation (**Table S2**). NRPS and PKS were particularly enriched in *Aspergillus* and *Fusarium*, two genera frequently linked to plant infections. Few BGC was identified in *Saccharomyces* and *Pichiomycetes*, consisting with existing research (Fig. 5a) [50, 51]. Then, we performed large-scale structure prediction of 386 representative core genes after clustering (DIAMOND threshold: 0.5) with Alphafold3 [52] to explore biosynthetic enzymes’ structure patterns in fungi. The results showed that core genes’ structures are highly diverse, but 21 NRPS-PKS hybrid enzyme clusters share high structural similarity (average pairwise TM-score > 0.50). By extracting these representative genes and mapping them to genomes, we discovered that NRPS-PKS enzymes in Fusarium are more conserved, while extensive sequence variations exist in Aspergillus (**Supplementary Fig. 6**).

To further validate BGC-Finder’s prediction, we focused on BGCs from *Aspergillus oryzae*, a well-studied model organism [53, 54]. We selected a NRP BGC containing one additional core gene, termed *Gene 363*, which was uniquely identified by BGC-Finder within the *AO-NSPID1 cluster 8.1* (Fig. 5b). A homolog search identified an isoprenoid synthase [55, 56] with high structure similarity (TM-score: 0.96) to *Gene 363.* For experimental validation, we knocked out *Gene 363* in *A. oryzae* and compared wild-type (WT) and mutant strains, confirming its functional role as a core gene. PCR analysis show that mutant strains were positive for 5F (∼1,400 bp) and 3F (∼1,650 bp) fragments (absent in WT) and negative for the RT (∼850 bp) fragment (present in WT), verifying successful disruption of *Gene 363* in *AO-NSPID1 cluster 8.1* (**Supplementary Fig. 7, 8**). Liquid chromatography-mass spectrometry (LC-MS) analysis of organic extracts from WT and mutant cultures revealed WT-specific peaks at m/z 379.2486, indicating *Gene 363* contributes to the biosynthesis of specific products. These differences were observed across different types of culture media (Fig. 5b for rice medium, **Supplementary Fig. 9** for CYP20 medium). The distinct metabolic profiles between WT and mutant strains confirm *Gene 363*’s biosynthetic role. Though novel BGCs may be particularly prevalent among uncultivated microorganisms, our findings suggest they may also be overlooked in well-characterized species.

### A plant infection-associated NRPS in two evolutionarily distant fungi

From seventeen *Fusarium* genomes, a diverse genus associated with plant and microbial interactions with significant implications for plant and food security, we identified 862 core genes within 625 BGCs. Phytotoxin synthases, notably AM-toxin (AMT) and HC-toxin (HCT) synthases, were significantly enriched in *Fusarium* **(Supplementary Fig. 10)**. We analyzed it at three different scales: (1) biosynthetic-core gene similarity, (2) BGC similarity, and (3) genus evolution timeline.

To expand putative phytotoxin BGCs, we compared known AMT and HCT core genes to all 862 core genes [36, 57], and identified 115 core genes which share sequence similarity. Given the modular nature of NRPS genes [58–61], we then mapped these core genes to their respective BGCs and clustered them into 59 GCCs (gene cluster clans) using BiG-SCAPE [36], including three AMT-related GCCs (GCC 540, GCC 1058, GCC 1703) and four HCT-related GCCs (GCC 21, GCC 101, GCC 792, GCC 455). Phytotoxin-related GCCs were predominantly enrich in *Fusarium*, and two HCT-related GCCs were found exclusively in *Coccidioides* (**Supplementary Fig. 10a**, Fig. 6a). GCC 1058 and GCC 1703, two AMT-related GCCs shared both BGC similarity and core gene structural similarity (**Supplementary Fig. 10b**, Fig. 6a). Furthermore, GCC 540, an AMT-related BGC family in *Fusarium*, exhibited substantial similarity with GCC 12, a siderophore-related BGC family in *Aspergillus* (Fig. 6a, b).

**Figure 6.**
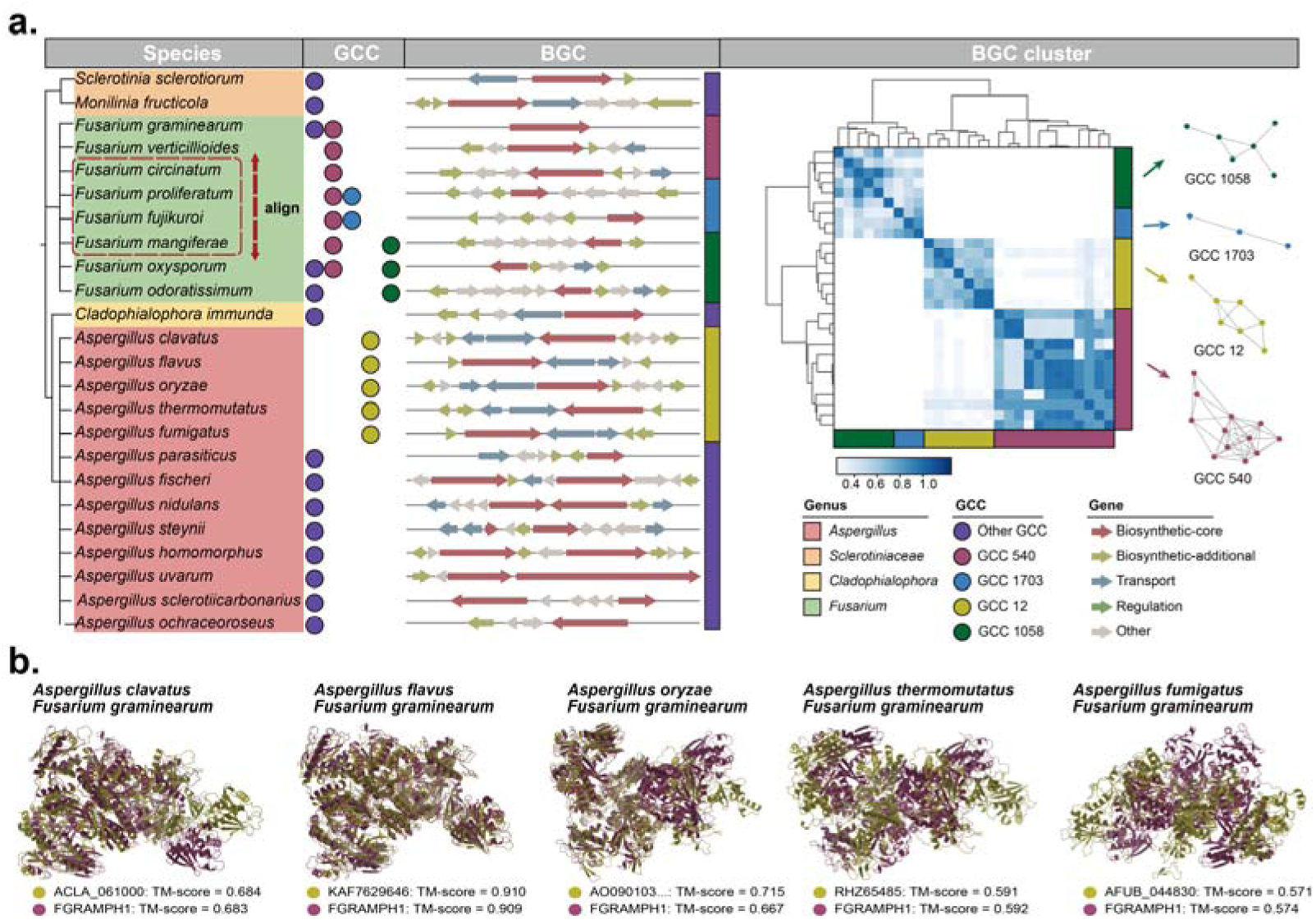
Evolutionary conservation of a phytotoxin-associated NRPS family in Aspergillus and Fusarium. **a.** Phylogenetic reconstruction of AM-toxin BGCs using reference AM-toxin synthase core genes aligned against FungiDB core genes, revealing three gene cluster clades (GCCs): GCC 540 (shared with Aspergillus siderophore BGC GCC 12), GCC 1058, and GCC 1703; adjacent cluster map illustrates BGC similarity. **b.** Structural comparison (TM-align) of NRPS core proteins from Fusarium GCC 540 (AM-toxin) and Aspergillus GCC 12 (siderophore).

To further investigate whether this similarity reflect horizontal gene transfer [62–64] or convergent evolution of NRPS BGCs in *Fusarium* and *Aspergillus*, we analyzed the evolutionary timeline and structural similarity of NRPS in these two genera. The evolutionary timeline indicates that these two genera diverged approximately 352 million years ago during the Paleozoic, an Era marked by extensive forest coverage. Despite this ancient divergence, their NRPS display remarkable structural similarity, a feature not observed in other co-evolving fungal species (**Supplementary Fig. 11**). We inferred that this NRPS family likely existed before their divergence, with both genera retaining it for synthesizing plant infection-related metabolites, while other species lost it over time, possibly due to their reliance on other fungal relatives [51, 65] (**Supplementary Fig. 12**). We inferred that differences in genomic context [66–68] and substrate specificity [69], driven by sequence variations, result in divergent product profiles [59, 70].

Our analysis has unveiled a diverse spectrum of phytotoxin BGC and suggests the evolution pattern of a NRPS family within two evolutionarily distant fungi: *Aspergillus* and *Fusarium* [50, 51, 71]. This finding enhances our understanding of fungal pathogenicity and offers insights for broad spectrum agricultural interventions targeting these biosynthetic pathways.

## Discussion

Traditional *in silico* BGC prediction methods suffer from long-stand limitations including time-consuming sequence alignment, manually defined-rules, *etc*. In this study, we presented BGC-Finder, a context-aware deep learning method that integrates protein language models with genomic information to predict and annotate BGCs. BGC-Finder leverages multi-scale architectures and attention mechanisms to capture both global genomic patterns and local sequence features, enabling the accurate and interpretable resolution of BGCs. It comprised of four models: two ESM2 models of different sizes to embed genes, a BGC-detector model to detect BGCs and an BGC-annotator model to annotate BGCs. We demonstrated that protein representations ESM2 models learned from pretraining stage enable function characterization, and the BGC-detector extracted latent biosynthetic patterns from microbial genomes. The BGC-annotator further leveraged bidirectional architecture to predict BGC class and gene function. Taken together, BGC-Finder achieves SOTA performance across all three tasks: BGC boundary detection, BGC class prediction and BGC gene annotation. The alignment-free nature of this pipeline accelerates BGC profiling speed up to 10^2^-fold compared to current approach.

Our interpretable analysis revealed that BGC-Finder autonomously captures the essential functional hierarchy of BGC genes through its attention mechanism without explicit biological priors. Crucially, the unexpected attribution of high importance to transporter genes aligns with recent research evidence [38], underscoring the model’s capacity to recover underappreciated biological principles *de novo*. We also discovered that BGC-Finder’s embeddings enable more cluster-contextualized representations than ESM2, and we utilized it to represent and group BGCs in parallel with annotation as an alternative to alignment-based approach. These results confirm that a supervised, pLM-based model can effectively learn genomic features from a modest amount of training data without large pretrained DNA language models like DNABERT [72] or Evo [73].

Additionally, we mined Arctic-Antarctic marine microbiome and discovered 1.2 times more BGCs which were missed by the alignment-based method. The microbial resources from geologically isolated biomes are unexplored and its biosynthetic potential is hidden by sequence divergence. Targeting at Arctic-Antarctic marine microbiome, we prioritized remote homologs, and we grouped BGCs and chose the main clusters containing BGCs only identified by BGC-Finder. There were 31 core biosynthetic genes in subclusters only showing structure similarity with homologs, demonstrating that BGC-Finder could uncover “microbial biosynthesis dark matter” by capturing shared structure information hidden in biosynthetic genes.

Furthermore, we conducted a comprehensive profiling of BGCs from fungal genomes and revealed the enrichment of NRPS and PKS in *Fusarium* and *Aspergillus*, two genera of significant ecological and agricultural importance. An NRPS gene family was discovered in both of them. We inferred that it existed before the divergence of them, and was evolved into distinct gene clusters and lay different functional roles. This finding provides new insights into the origin and evolution of NRPS BGCs.

Our study also has several limitations. The transformer-based models require high-memory GPUs to accelerate inference, which limits BGC-Finder’s accessibility. To mitigate this, we developed an online, user-friendly BGC-Finder pipeline and released it on Google Colab. Future work could focus on optimizing computational efficiency to enhance accessibility. Besides, we refer to ‘microbial biosynthesis dark matter’ as biosynthetic genes cannot be detected by sequence alignment, rather than completely novel or unknown class of BGCs like ‘thiopeptide synthase’ and ‘arylpolyene synthase’ in this article. Introducing an enzyme annotation module into the pipeline may help to cover novel class of BGCs. Last but not least, in microorganisms, especially fungi, some enzymes involved in the synthesis of secondary metabolites aren’t co-localized in the genome, and these synthesis pathways are difficult to capture with existing methods. It is anticipated that the introduction of a metabolic pathway annotation module into our pipeline in the future to solve this problem.

In conclusion, BGC-Finder enables accurate, rapid, and interpretable BGC profiling. It replaces fragmented profiling pipelines with an end-to-end system that detects, annotates and clusters BGC, enabling large-scale biosynthetic discovery while illuminating “microbial biosynthesis dark matter” that were untapped by other methods. Its application to fungi and bacteria genomes has provided novel insights and experimentally validated discoveries, demonstrating its potential to advance our understanding of microbial secondary metabolism and the discovery of ‘microbial biosynthesis dark matter’. Furthermore, by elucidating the functional roles of genes within BGCs, BGC-Finder also paves the way for the design and engineering of bespoke biosynthetic pathways.

## Code availability

The source code for BGC-Finder is available at https://github.com/HUST-NingKang-Lab/BGC-Finder. The Colab BGC-Finder is available online: https://colab.research. google.com/drive/1ToMM3foB77ssSjHjZ43cBhXcKFD6k9SZ?usp=sharing. Its step-by-step tutorial is available at https://github.com/HUST-NingKang-Lab/BGC-Finder /blob/main/BGC-finder_step_by_step.ipynb.

## Methods

### Data collection

We collected three datasets in our study: data for training, data for validation and data for mining.

#### Data for training

A total of 2,499 BGCs from MIBiG Version 3.1 were collected. Each sample was labeled with both product type and gene function. BGCs with multiple products were duplicated and labeled with a single product to ensure each sample was associated with only one product, resulting in 2,994 samples. Of these, 2,394 BGCs were allocated for model training (MIBiG training-set), 300 for internal validation (MIBiG validation-set), and 300 for model test (MIBiG test-set).

#### Date for validation

The nine bacteria genomes containing 291 annotated biosynthetic gene clusters (BGCs) from Cimermancic et *al.* [12] were utilized for model evaluation with DeepBGC and BGC-Prophet. Additionally, FunBGCs, a manually curated dataset of fungal BGCs, [20] served as a benchmark in core gene discovery task.

#### Data for mining

A total of 210 metagenome-assembled genomes (MAGs) were collected from the research in Arctic and Antarctic marine microbiota [74]. The taxonomy annotation was performed using the module “classify_wf” of Genome Taxonomy Database Toolkit (GTDB-Tk, version 2.2.6) against the GTDB with default parameters. A total of 256 genomes representing 197 fungal species from FungiDB latest version (68), as reported by Basenko et al. [18], were analyzed to identify putative fungal BGCs and biosynthetic core genes. Each gene in the genomes was annotated with Gene Ontology (GO) terms, and the results obtained from BGC-Finder were cross-referenced with these annotations to identify additional potential core genes.

### Establishment of BGC-Finder

#### BGC-detector model architecture

The BGC identification model is a transformer-based model we have developed and published: BGC-Prophet. It utilized ESM2-8M to encode each gene in the genome into a 320-dimensional embedding, and then identifies BGC genes in a context length of 128.

#### BGC-annotator model architecture

We developed the BGC-annotator model based on RoBERTa architecture aligned with gLM. BGC-Finder consisted of 5 layers, each with a hidden size of 1280 and 10 attention heads per layer, utilizing relative positional embeddings (‘relative_key_query’) [75]. Each transformer layer comprises of a self-attention layer and a fully connected layer, allowing the model to capture complex dependencies within genomic data. The classification head is implemented as a single-layer feed-forward neural network, with an output layer dimension that corresponds to the number of gene and class categories to be predicted. Specifically, the gene categories include: Biosynthetic-core (core), Biosynthetic-additional (tailoring), Transport, Regulation, and Other. The class categories consist of: Non-Ribosomal Peptide (NRP), Ribosomally Synthesized and Post-Translationally Modified Peptide (RiPP), Alkaloid, Polyketide, Saccharide, Terpene, and Other.

Each gene in the samples was characterized by a 1280-dimensional embedding generated by ESM2-650M, and mean pooling was applied to the token embeddings to generate a single, fixed-length vector representation for each gene. Class embeddings were initialized with zero as a CLS token, serving as a representative embedding for class prediction and concatenated with the gene embeddings to form a unified representation of the BGCs. To distinguish the class token and gene token, the ‘token_type_id’ of class is set to 0 and the ‘token_type_id ’of genes is set to 1. Given the fixed input length requirement of transformer models, we set input length to 128 to ensure adequate coverage of BGC size and each position in the sequence is represented by a 1280-dimensional vector. The input representation for a single BGC is formally defined as:

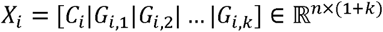

where *X_i_* denotes a single BGC, *C_i_* represents the class embedding vector, *G_i,j_* ∈ ℝ*^n^*^×1^ represents the embedding vector of the j-th gene in the BGC embedding, n = 1280 is the dimensionality of each embedding vector and k is the number of genes in the BGC, with padding or truncation applied as needed to match the fixed input length. Before input to the transformer block, X was added to the relative position embeddings and token type embeddings.

The self-attention mechanism used in RoBERTa adheres to the scaled dot-product attention formulation:

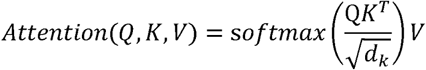

where Q (queries), K (keys), and V (values) are linear projections of the input, and *d_k_* is the dimensionality of the keys. This formulation allows the model to capture contextual relationships among genes and product embedding within the sequence.

#### Training details

The Adam optimizer was applied with an initial learning rate of 5 X 10^-5^. The model parameters were initialized with gLM. Training was set to run for a maximum of 100 epochs, employing an early stopping strategy to retain the best-performing model. For hyperparameter selection, we conducted a grid search in the internal validation-set and selected a structure comprising 5 layers and 10 attention heads (**Supplementary Fig. 1**). The modeling framework was implemented in PyTorch, utilizing the HuggingFace Transformers library for model configuration and training.

#### Generation of context-aware embeddings

To generate global landscape of BGC embeddings extracted from BGC-Finder, all 204,661 BGCs were downloaded from BiG-FAM database, and 23,000 representative BGCs from 23 main categories were downloaded for analysis. The context-aware embeddings generated by the model were extracted from each layer, including a word embedding layer and five encoder layers. The embeddings of gene token and class token were averaged to keep context information. To get context-aware embeddings generated by ESM2, each gene’s embedding was averaged.

### Pipeline of BGC-Finder

#### Gene prediction and BGC screening

Prodigal v2.6.3 [32] was used to predict genes from bacteria genomes. AUGUSTUS [33] was used to predict genes from fungi genomes. Before inference, genes were grouped into batches based on their genomic location, including metagenome contig number or chromosome number. Each gene was encoded by ESM2-6M as an embedding of length 320, and BGC genes were screened using BGC-Prophet. Genes with probability > 0.5 were selected as candidate BGC genes.

#### BGC annotation and reconstruction

Each BGC gene was encoded by ESM2-650M as an embedding of length 1280. The annotation model was employed to predict each gene function and BGC class. The protein sequences and corresponding DNA of all BGCs were retrieved from genomes and reconstructed into GenBank format files using Python Package Biopython version 1.83.

### Model evaluation

To assess classification performance of BGC-Finder, we computed areas under the receiver operating characteristic curve (AUROC) and areas under Precision-Recall curve (AUPR) for both gene classification and product prediction tasks using MIBiG test-set. The ROC curve and AUROC value were calculated using scikit-learn methods. The RF was retrained separately for gene classification and product prediction, and ROC curve and AUROC score were calculated separately to assess its individual performance on each task. In the external benchmark, we compared the performance of different methods for their ability to accurately detect BGC genes. The evaluation metrics included accuracy, F_1_-score, precision and recall:

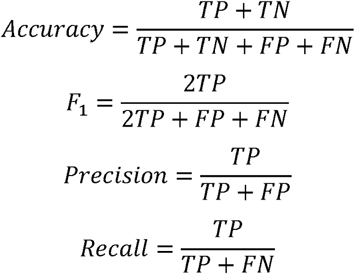

where TP, FP, TN and FN are the number of true positives, false positives, true negatives and false negatives predicted, respectively.

To assess protein encoding strategy and model architecture, we further trained two Random Forest (RF) models, RF-Gene for gene function annotation task and RF-Production product prediction task, on the same model training-set. Models based on ESM2 embeddings significantly outperformed models based on one-hot encodings in both tasks (**Supplementary Fig. 3**). Notably, ESM2 embeddings could capture key biosynthetic genes’ features by placing them into more similar space (P < 0.0001) (**Supplementary Fig. 4**). While RF-Gene matched BGC-Finder’ s performance in gene function prediction, RF-Product’s average AUROC for product classification was only 0.791, even with ESM2 embeddings. This disparity underscores the advantage of BGC-Finder’s transformer-based architecture over traditional machine learning models in leveraging genomic features for product inference.

### Model interpretability

To assess the interpretability of the BGC-Finder model and its ability to capture core gene features, we focused on analyzing the model’s attention mechanisms. Specifically, we extracted attention weights from all attention heads (n=50) by running inference on all sequences in the MIBiG test set. To evaluate the model’s capacity to capture core gene features, each sequence from the test set was represented as a binary vector of BGC length, where a value of 0 indicates a non-core gene and 1 indicates a core gene. We calculated the Spearman correlation between the attention weights and the presence of core genes using the scikit-learn library. We further explored the gene contribution to product inference by examining the attention weights between gene embeddings and product embeddings. Specifically, attention weights were extracted where product embeddings served as Queries (Q) and gene embeddings acted as Keys (K), and these attention scores are referred to as gene-product scores. To highlight the roles of different genes in product inference, the top 5% of gene-product scores from each gene category were collected and displayed.

### BGC clustering

BiG-SCAPE [36] was employed to analyze the Gene Cluster Families (GCFs) and Gene Cluster Contexts (GCCs) for candidate BGCs. DSS value (Domain Sequence Similarity) was used to measure the similarity between two BGCs as follows:

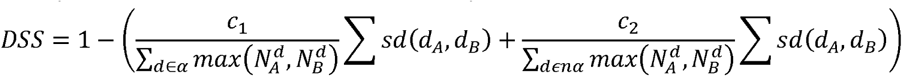

Where a represents the anchor domains, a list of domains can be given a special weight, such as those in NRPS or PKS, and na accounts for the remaining domains; d are all distinct domain types in the BGC pair, and N^d^ donates the number of copies of domain d in the BGC. sd computes the sequence dissimilarity between domain d in BGC A and B by taking all copies of the domain and summing the complement of their sequence identities. c_1_ and c_2_ are weighting coefficients that adjust the contributions of anchor and non-anchor domains, respectively.

### Sequence alignment and clustering

DIAMOND [76] with default parameters was used to align sequences and sequence clustering. To search homologs in the public database, Foldseek server [42] was used to query sequences, and sequences shared highest sequence identity with function annotation were selected for further analysis.

### Calculation of sequence diversity

To calculate sequence diversity, all sequences were firstly clustered by DIAMOND at threshold of 0.9. Then sequence diversity was calculated as follows:

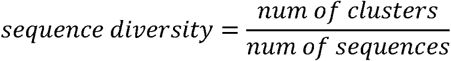

### Structure prediction and alignment

For protein structure prediction, AlphaFold3 [52] generated the tertiary structures of selected proteins. We used TM-align [77] to compare and align protein structures, computing the TM-score to evaluate structural similarity as follows:

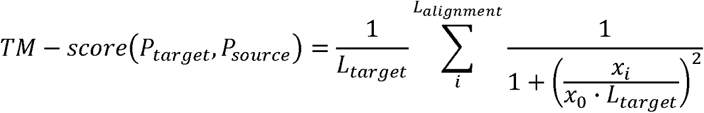

Where *P_source_* is the source protein aligned to target protein *P_target_. L_target_* is the length of target protein, and *L_alignment_* is the number of paired residues. *x_i_* is the distance between the i-th paired residues. All procedures followed default settings.

### DNA extraction

*Aspergillus oryzae* NSPID1 (AO-NSPID1) was obtained from our laboratory for the following gene deletion and validation. AO-NSPID1 glycerol stock was retrieved from -80°C and thawed. A 40 μL aliquot was transferred onto a PDA + UU agar plate using a sterile pipette tip. The sample was evenly spread across the surface with a sterile disposable spreader, and the plate was sealed, labeled, and inverted for incubation at 30°C for 5 days. Fungal biomass was dried and transferred to a 2 mL tube with 400 μL LETS Buffer and 3 steel beads, then homogenized at 70 Hz for 60 seconds. After a 5-minute rest, 300 μL LETS Buffer was added, followed by a 5-minute incubation. PCI extraction was performed by adding 700 μL of the lower phase, shaking for 20 seconds, and centrifuging at 13,000 rpm for 10 minutes at 4°C. The aqueous phase was transferred to a new tube, mixed with 1 mL pre-cooled 95% ethanol, and centrifuged at 13,000 rpm for 10 minutes. The pellet was washed with 300 μL 75% ethanol, air-dried, and DNA was eluted with 30 μL ddH2O at 65°C. The genomic DNA was quantified, diluted to 50–100 ng/μL, and stored at -20°C.

### Gene amplification

#### PCR procedure

Recombinant plasmids were constructed using SnapGene software to insert the target gene into a suitable plasmid. Primers for each fragment were designed, including forward (FW) and reverse (RV) primers. The primers were approximately 20 bp in length, with an annealing temperature of around 55°C and homologous arm lengths of 20 bp. The PCR reaction mixture was prepared with the final addition of 2× Phanta Flash Master Mix. The Master Mix was stored at -20°C after use. The PCR amplification program began with pre-denaturation at 98°C for 10 seconds, followed by 34 cycles of denaturation at 98°C for 10 seconds, annealing at the primer-specific melting temperature for 5 seconds, and extension at 72°C for 5 seconds per kilobase. A final extension at 72°C for 2 minutes completed the process, with the products stored at 4°C.

#### DNA Agarose Gel Electrophoresis

For DNA separation, 0.6 g of high-purity, low electro-osmotic agarose was mixed with 60 mL of TAE buffer (30 mL for smaller volumes). The solution was microwaved for 2 minutes to dissolve, followed by an additional 30 seconds of heating until visible bubbles formed. After slight cooling, a red fluorescent DNA dye was added. The agarose solution was poured into the gel mold, and combs were inserted to create wells. The gel was allowed to solidify for approximately 20 minutes, after which the combs were carefully removed. The gel was placed into an electrophoresis chamber, ensuring that the TAE buffer completely covered the gel. The wells were oriented towards the negative electrode. PCR products were mixed with 6× Loading Buffer, and 30 μL of the mixture was loaded into each well along with 5 μL of DL 5000 DNA Marker. Electrophoresis was performed at 120 V (not exceeding 140 V, 200 mA) for approximately 25 minutes. The gel was then visualized under UV light using a gel imaging system, with the image captured after adjusting the brightness and contrast for clarity.

### Gene knockout experiment

#### Fungal preparation

The germinated Aspergillus oryzae culture was filtered through an iron sieve, washed with 200 mL of water, pressed dry, and transferred to a 50 mL centrifuge tube. Yatalase (120 mg) was dissolved in 20 mL of Solution 0 (final concentration: 60 mg/mL), filtered through a 0.22 μm membrane, and added to the fungal biomass. The mixture was incubated at 30°C and 120 rpm for approximately 60 minutes, with protoplast formation monitored at 50 minutes. After 100 minutes, the suspension was filtered through a glass funnel, kept on ice, and centrifuged at 1000 g, 4°C, for 5 minutes. The supernatant was discarded, and the pellet was gently resuspended in 10 mL of 0.8 M NaCl. Protoplasts were counted, with each experimental group containing 4 × 10L protoplasts. The suspension was centrifuged at 900 g, 4°C, for 5 minutes, the supernatant was discarded, and the pellet was resuspended in Solution II and III (4:1, 100 μL per group).

#### Fungal transformation

The DNA solution was prepared by dissolving at least 10 μg of DNA in 50 μL of sterile water and mixing with 50 μL of 2× Solution II. A 100 μL aliquot of the protoplast suspension was added to the DNA solution and incubated on ice for 20 minutes. Solution III (50 μL) was then added, followed by incubation at 25°C for 20 minutes. Subsequently, 2 mL of Solution III was added in steps, gently mixed, and incubated at room temperature for 5 minutes. The mixture was then diluted with 4 mL of Solution II. A 600 μL aliquot of the transformed protoplast suspension was plated onto each CD plate, with 10 plates per group, and incubated at 30°C overnight. The following day, 5 mL of Top Agar was overlaid onto each plate. Once solidified, the plates were sealed and incubated upside down at 30°C. Transformant growth was assessed on the third day. To screen positive transformants, secondary screening plates (8 mL each) were prepared and transformants were picked using a sterile pipette tip, transferred to the plates, labeled, and incubated at 30°C.

#### Gene knockout validation

To obtain fungal biomass, MPY liquid medium was prepared, sterilized at 121°C for 30 minutes, and distributed into sterile 40 mL vials, each containing 5 mL. Biomass was acquired from the rescreening plates and added to the vials. A control was also prepared by adding wild-type Aspergillus oryzae spores to another vial. The vials were sealed with breathable sealing film, labeled, and incubated at 30°C and 220 rpm for 3-5 days. Genomic DNA was extracted from each transformant. Primers were designed for PCR validation of each knockout strain. For the 8.1-5F, 8.1-3F, and 8.1-RT fragments, the respective i5F-FW, i5F-RV, i3F-FW, i3F-RV, iRT-FW, and iRT-RV primers were used. PCR amplification was performed using the extracted genomic DNA as templates. The PCR program included initial denaturation at 98°C, followed by 34 cycles of denaturation, annealing, and extension, with final extension at 72°C. Validation of knockouts was confirmed based on the presence or absence of PCR amplicons.

### Fungi Fermentation

#### Transformant fermentation

CYA, CYA20, and YES small-scale fermentation was conducted [78]. The respective liquid media were prepared, sterilized at 121°C for 30 minutes, and distributed into sterile 40 mL vials, each containing 5 mL. Spore suspensions were added to the vials, with a control vial containing wild-type Aspergillus oryzae spores. The vials were sealed with breathable sealing film and incubated at 30°C and 220 rpm for 5 days.

#### Rice small-scale fermentation

For the rice small-scale fermentation, 2 g of rice and 1.8 mL of water were added to 40 mL vials, sealed with breathable sealing film, and sterilized at 121°C for 30 minutes. After cooling, spore suspensions were added to the vials, with a control vial containing wild-type Aspergillus oryzae spores. The rice was loosened with a sterile pipette to ensure homogeneity. The vials were sealed with breathable sealing film and incubated at 30°C for 12 days.

#### Extraction of fermentation products

Fermentation products from liquid and solid media were extracted using industrial-grade methanol overnight. The methanol extracts were concentrated by rotary evaporation, reconstituted in chromatographic-grade methanol, and stored at -20°C.

### LC-MS Analysis of Fermentation Products

LC-MS analysis of crude extracts was performed using an Eclipse Plus C18 column (2.1 × 100 mm, 3.5 μm). The mobile phase consisted of CH_3_CN and H_2_O (both containing 0.1% HCOOH). The CH3CN concentration was increased linearly from 5% to 98% over 11 minutes and maintained at 100% for 2 minutes. The flow rate was 0.5 mL/min, and the column temperature was set at 25°C.

### Visualization of embeddings

We visualize BGC-Finder’s embedding using a uniform manifold approximation and projection for dimension reduction (UMAP). Using the Python package, umap [79], we transform BGC-Finder’s embeddings to two-dimensional vectors via the parameters: n_neighbors = 10, min_dist = 0.10, n_components = 2 and the cosine distance metric.

### Data visualization

The Python packages Matplotlib [80] and Seaborn [81] were used for data visualization, while iTOL [82] was employed for tree visualization.

## Acknowledge

This work was partially supported by the National Key R&D Program of China (Grant No. 2023YFA1800900, 2021YFA0910500 and 2018YFC0910502) and the National Natural Science Foundation of China (Grant Nos. 32071465, 31871334, 81827901, 22277035).

## Author contribution

K.N., Y.Z. and H.B. conceived and proposed the idea. Z.K., H.Z. and C.L. performed the experiments and analyzed the data. Z.K. and H.Z. visualized the data. Z.K., H.Z., Y.Y., Y.Z., K.N. and H.B. contributed to editing and proof-reading the manuscript. R.Y. joined the discussions. All authors read and approved the final manuscript.

## Supplementary

**Supplementary Figure 1.**
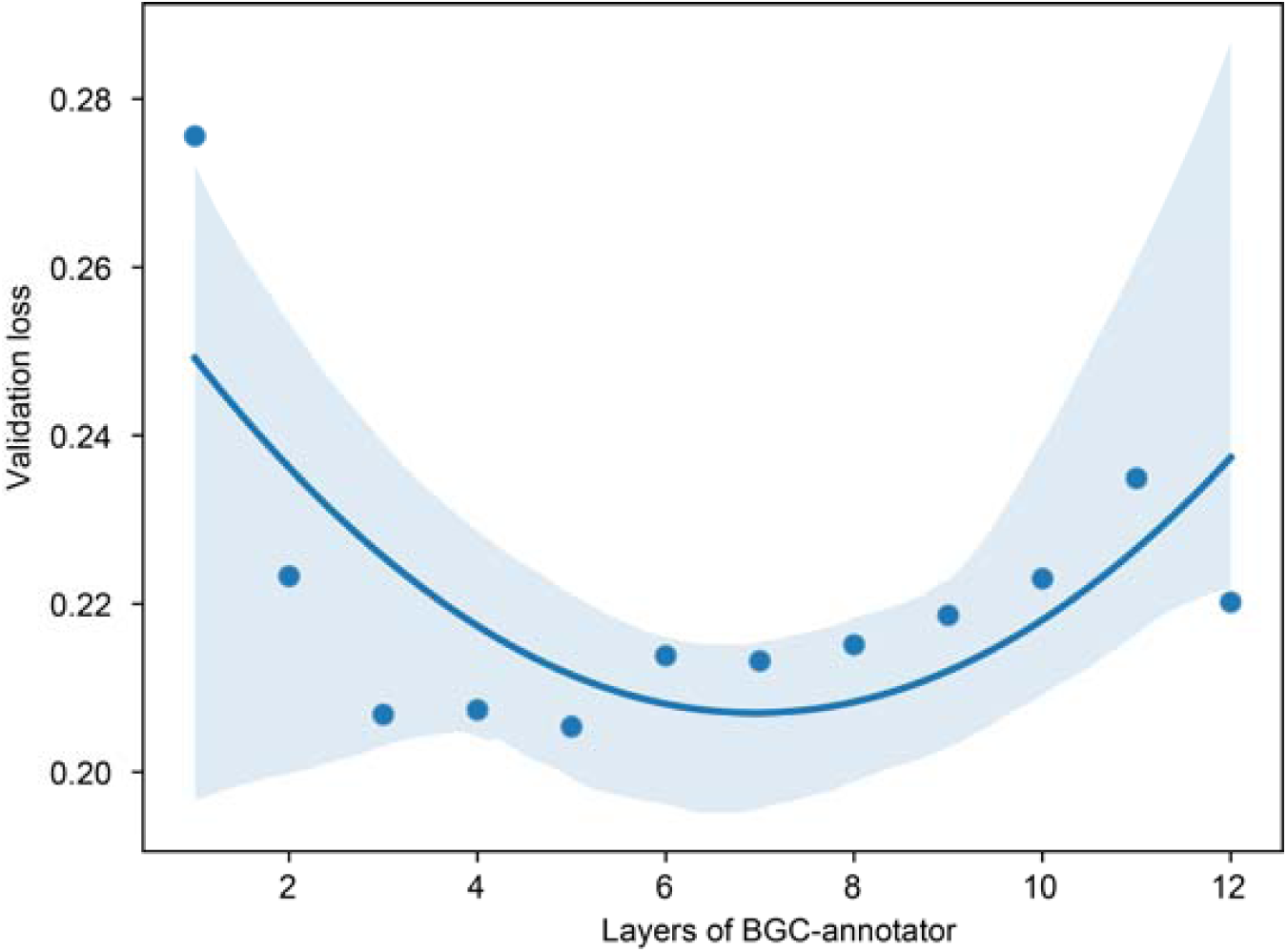
Validation loss across grid search configurations. Rows indicate the model’s validation loss, and columns represent the number of layers. The model achieved the lowest loss using five attention layers.

**Supplementary Figure 2.**
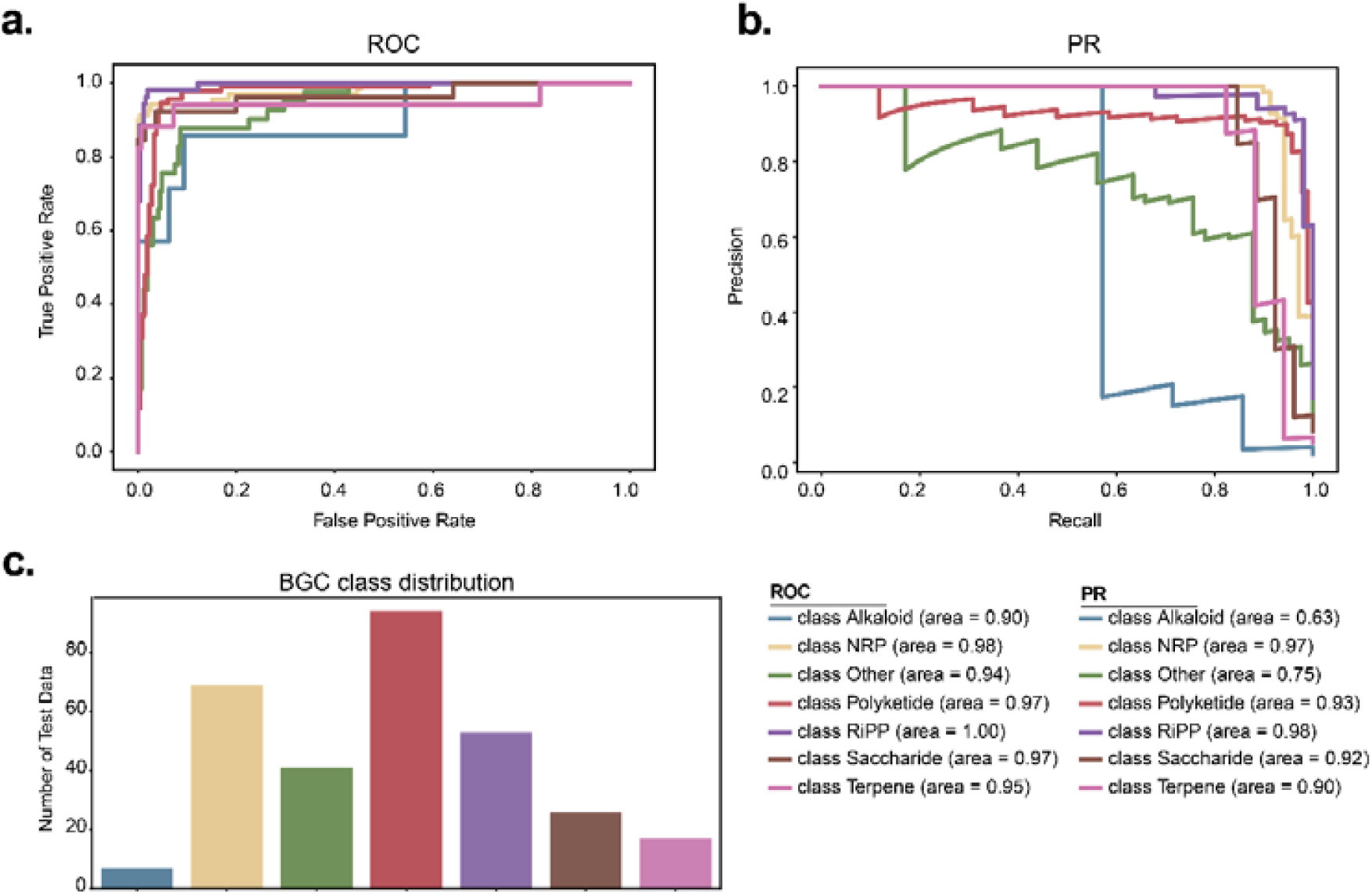
Performance of BGC-Finder on MIBiG test-set and data distribution. **a.** Precision-Recall curve for BGC gene detection by BGC-Finder.**b.** Precision-Recall curve for BGC class prediction by BGC-Finder. **c**. Data distribution of BGC classes in the MIBiG test set (e.g. Alkaloid: 7, Terpene: 17).

**Supplementary Figure 3.**
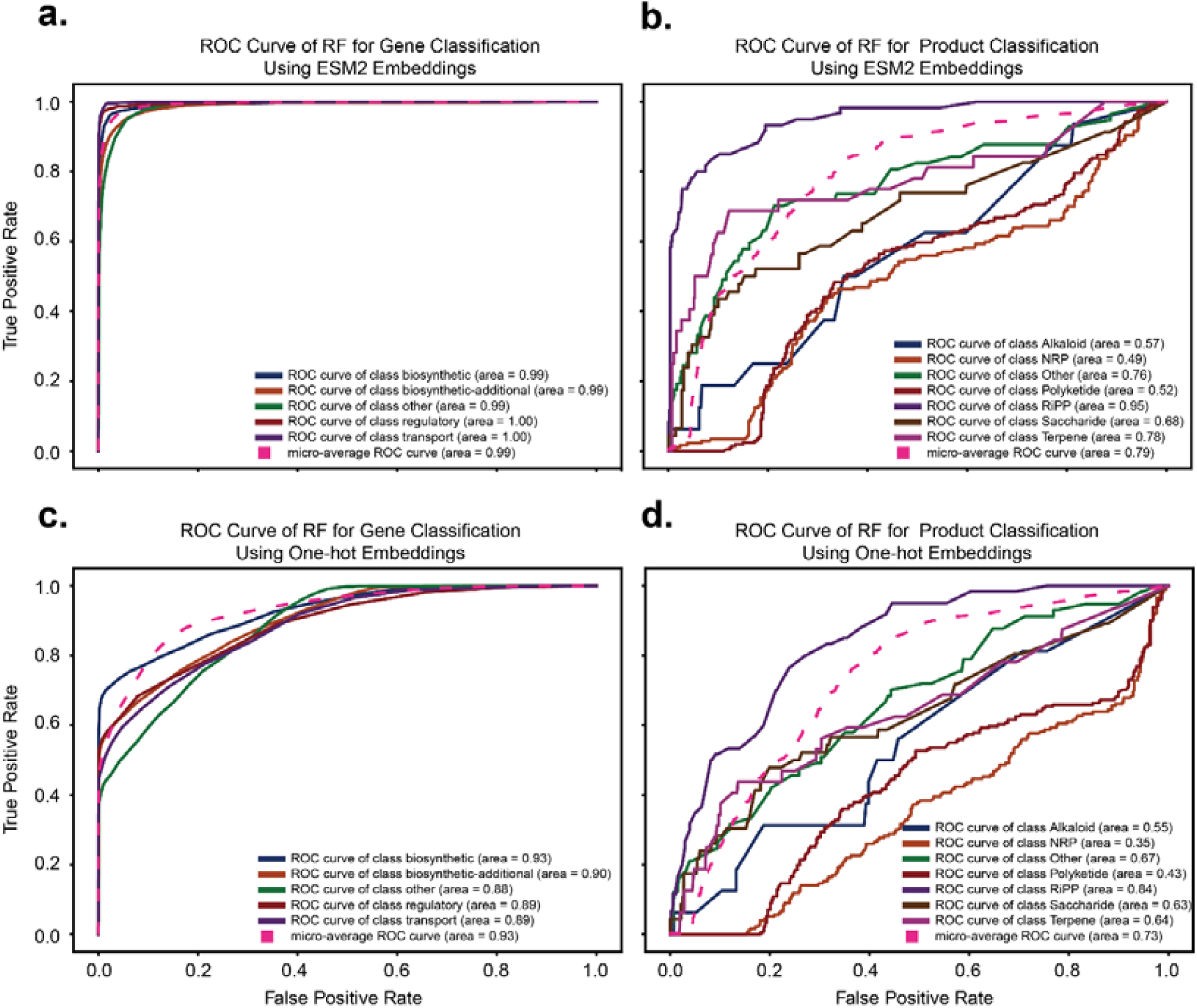
Random-Forest (RF) Classification Performance using ESM2 and One-Hot embeddings. **a.** ROC curve for RF gene classification in MIBiG test-set using ESM2 embeddings. **b.** ROC curve for RF product class classification in MIBiG test-set using ESM2 embeddings. **c.** ROC curve for RF gene classification in MIBiG test-set using One-hot embeddings. d. ROC curve for RF product class classification in MIBiG test-set using One-hot embeddings.

**Supplementary Figure 4.**
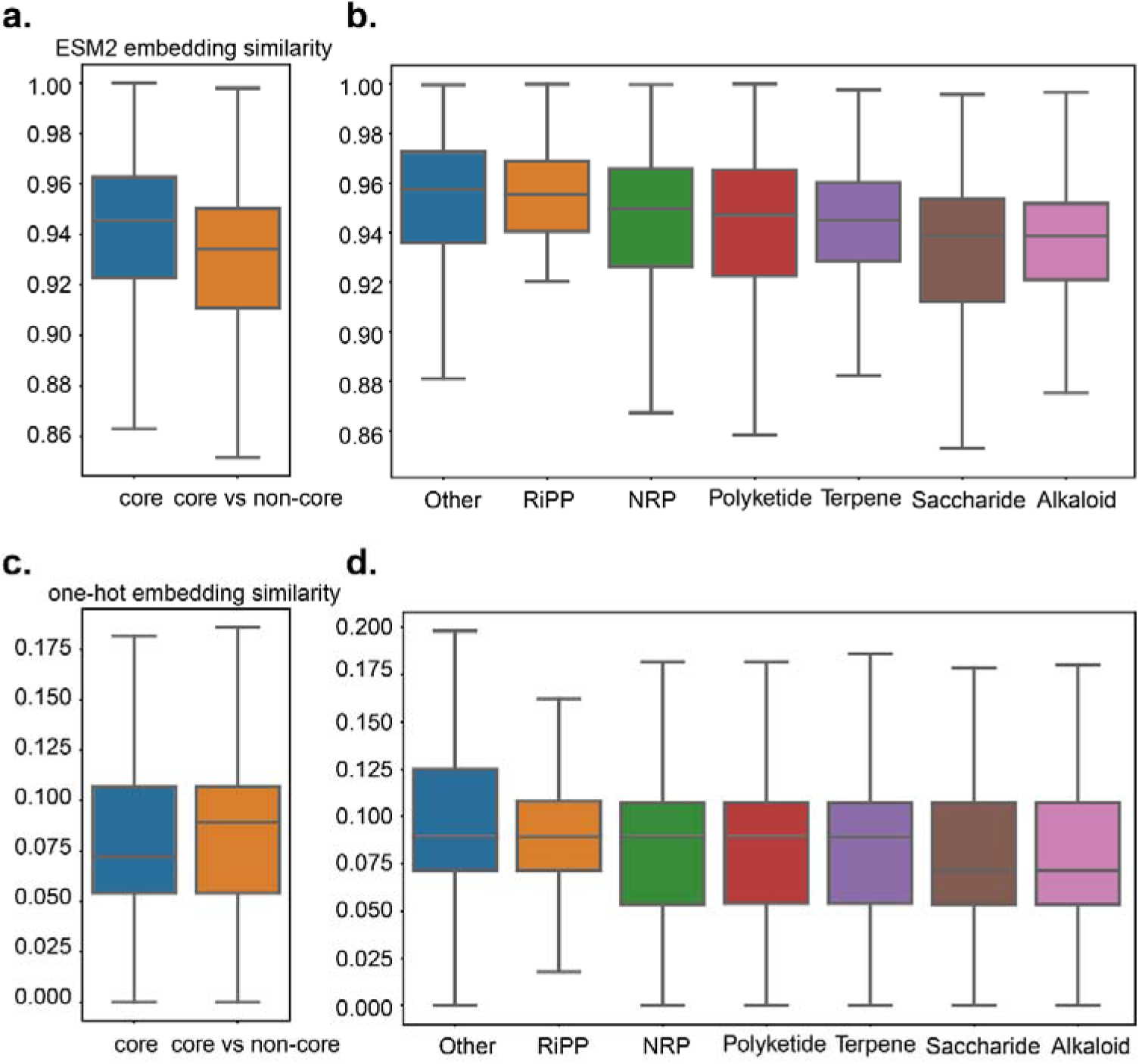
Cosine similarity across gene embeddings. **a**. Internal similarity of core genes and core vs. non-core gene similarity using ESM2 embeddings (p-value: 2.96e-48) **b.** Internal and global similarity of core genes with different products using ESM2 embeddings c. Internal similarity of core genes and core vs. non-core gene similarity using one-hot embeddings (p = 0.51) b. Internal and global similarity of core genes with different products using one-hot embeddings

**Supplementary Figure 5.**
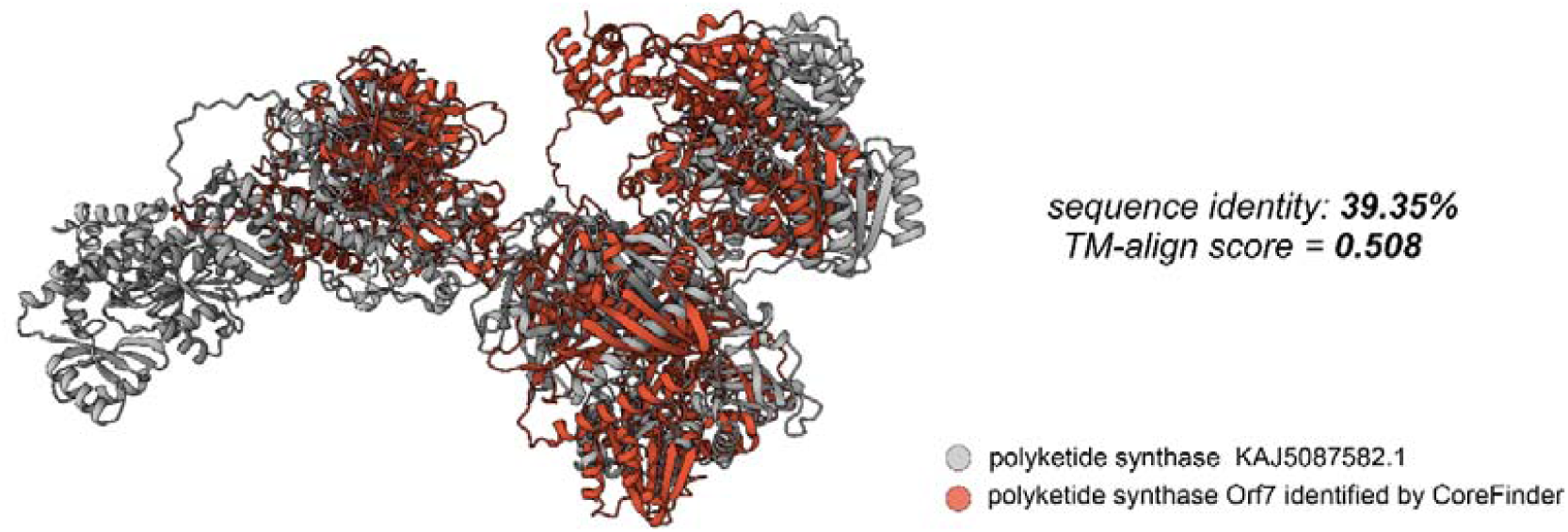
Protein structure visualization. The polyketide synthases identified by BGC-Finder was aligned with the experimentally verified polyketide synthase (KAJ5087582.1) structures with the highest sequence similarity (remote homolog). The polyketide synthase Orf7 was annotated as ‘other’ by antiSMASH.

**Supplementary Figure 6.**
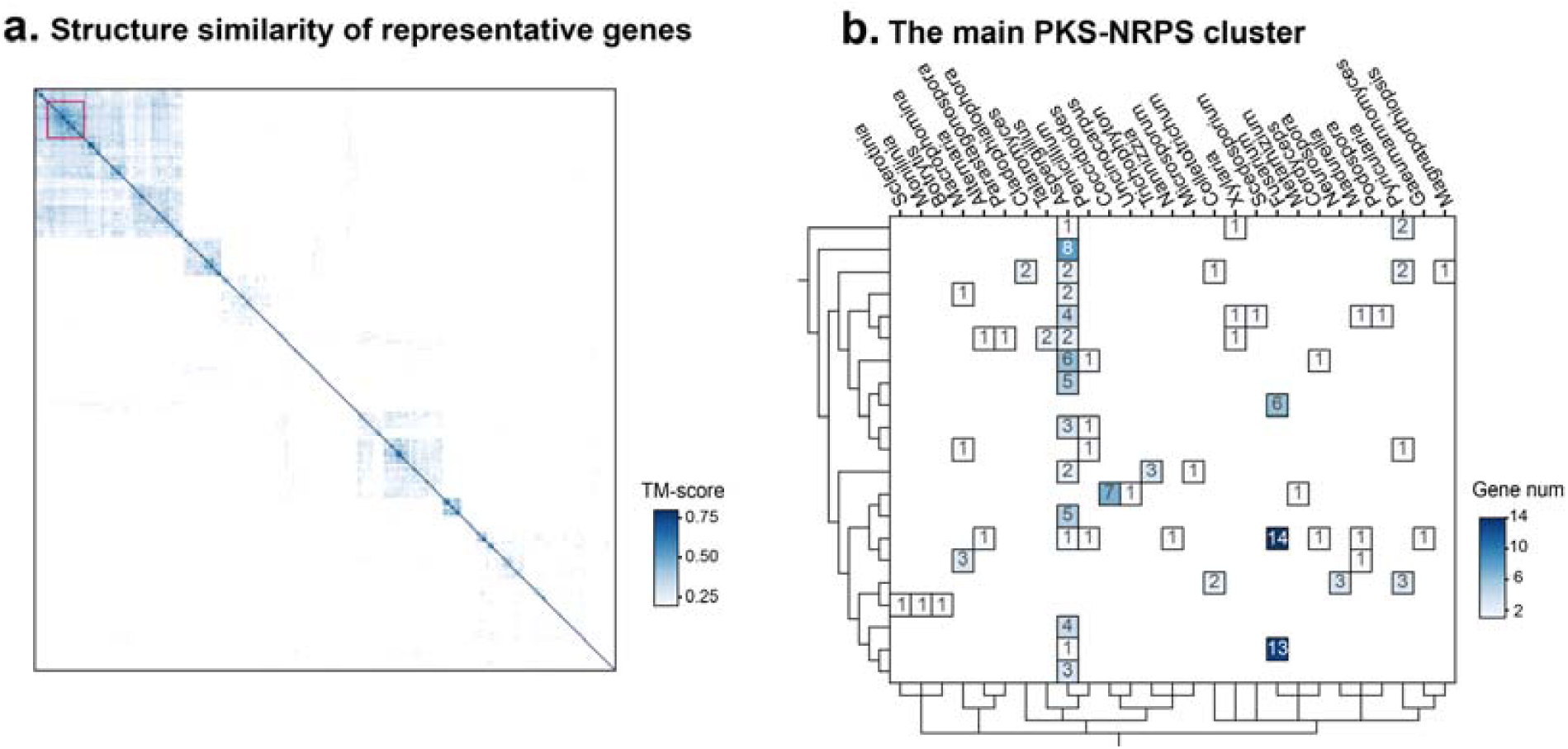
The Structure similarity of representative genes and main PKS-NRPS cluster shares high structure similarity. **a.** All biosynthetic-core genes identified from FungiDB were clustered using DIAMOND with threshold of 0.50. The representative genes from 396 clusters were extracted for structure prediction using Alphafold 3. Pair-wise TM-scores were calculated, and 21 representative genes related to PKS-NRPS shared high structure similarity. **b.** The order of the columns is based on phylogenetic information, and the order of rows is based on representative genes’ structure similarity.

**Supplementary Figure 7.**
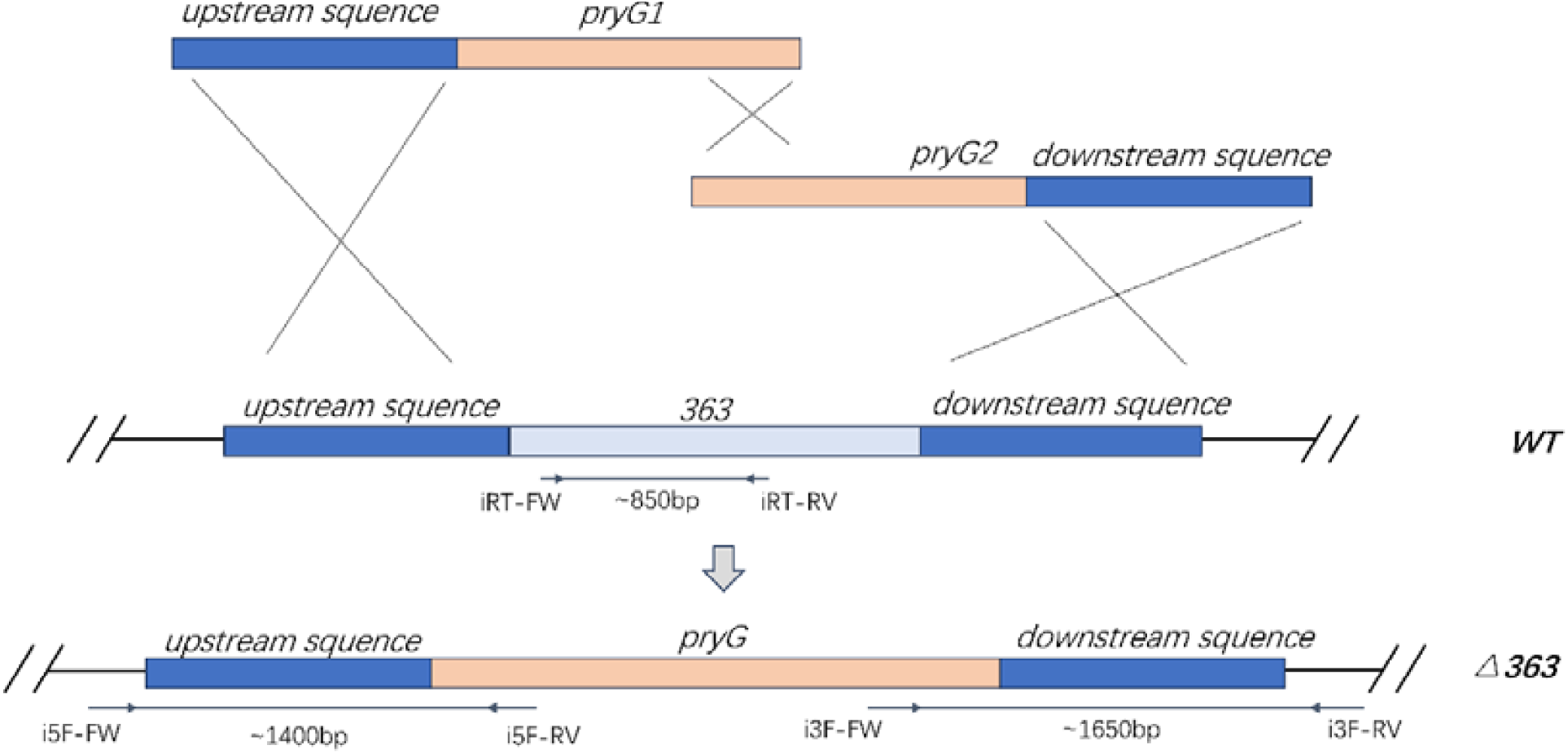
Schematic representation of the splitDmarker strategy used to delete the 363 gene (labeled “363” in the wild type, WT) and the corresponding PCR verification steps. In this approach, the upstream flanking region of *363* is fused to a partial selectable marker (*pryG1*), while the downstream flanking region is fused to a second partial selectable marker (*pryG2*). These two partial markers share overlapping sequences that reconstitute a functional *pryG* gene through homologous recombination, thereby replacing the *363* locus and generating the Δ*363* strain. To confirm the deletion, primers iRT-FW and iRT-RV were used to amplify an ∼850 bp internal fragment of *363* (present in the WT but absent in the Δ*363* mutant). Primer sets i5F-FW / i5F-RV (∼1400 bp) and i3F-FW / i3F-RV (∼1650 bp) were employed to validate the correct integration of the *pryG* replacement cassette at the 5′ and 3′ flanking sequences, respectively. Blue boxes indicate genomic regions flanking the *363* locus, orange boxes indicate partial selectable marker sequences (*pryG1* or *pryG2*), and the gray lines show homologous recombination events.

**Supplementary Figure 8.**
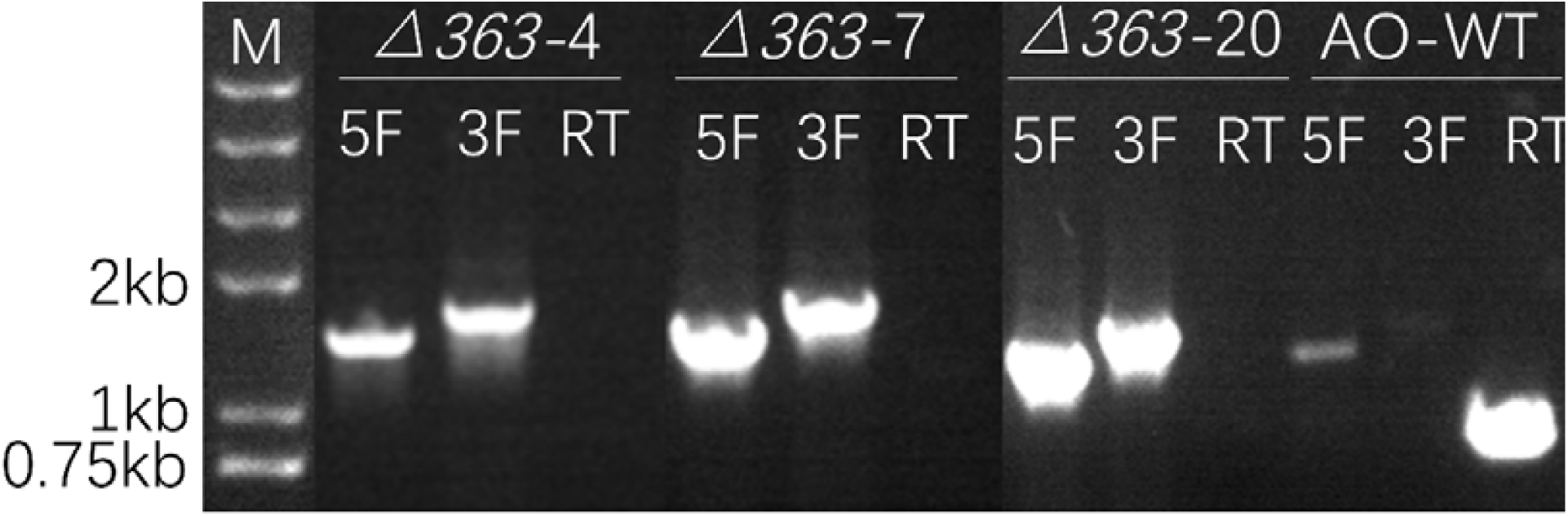
PCR verification of gene *363* knockout mutants generated via the split-marker strategy. Gel electrophoresis of PCR products from three representative mutants (Δ*363*-4, Δ*363*-7, Δ*363*-20) and the wild-type control strain (AO-WT). Primer sets labeled 5F and 3F were used to verify the correct integration at the 5’ and 3’ flanking regions, respectively. The internal RT-PCR assay (RT), amplifying a segment within the *363* gene, confirms successful gene disruption by the absence of the PCR product in mutants. Mutants Δ*363*-4, Δ*363*-7, and Δ*363*-20 exhibit the expected amplified bands at approximately 1.4 kb (5F) and 1.65 kb (3F), while the wild-type strain (AO-WT) only yields the internal 363 gene band (RT). DNA marker sizes are indicated on the left.

**Supplementary Figure 9.**
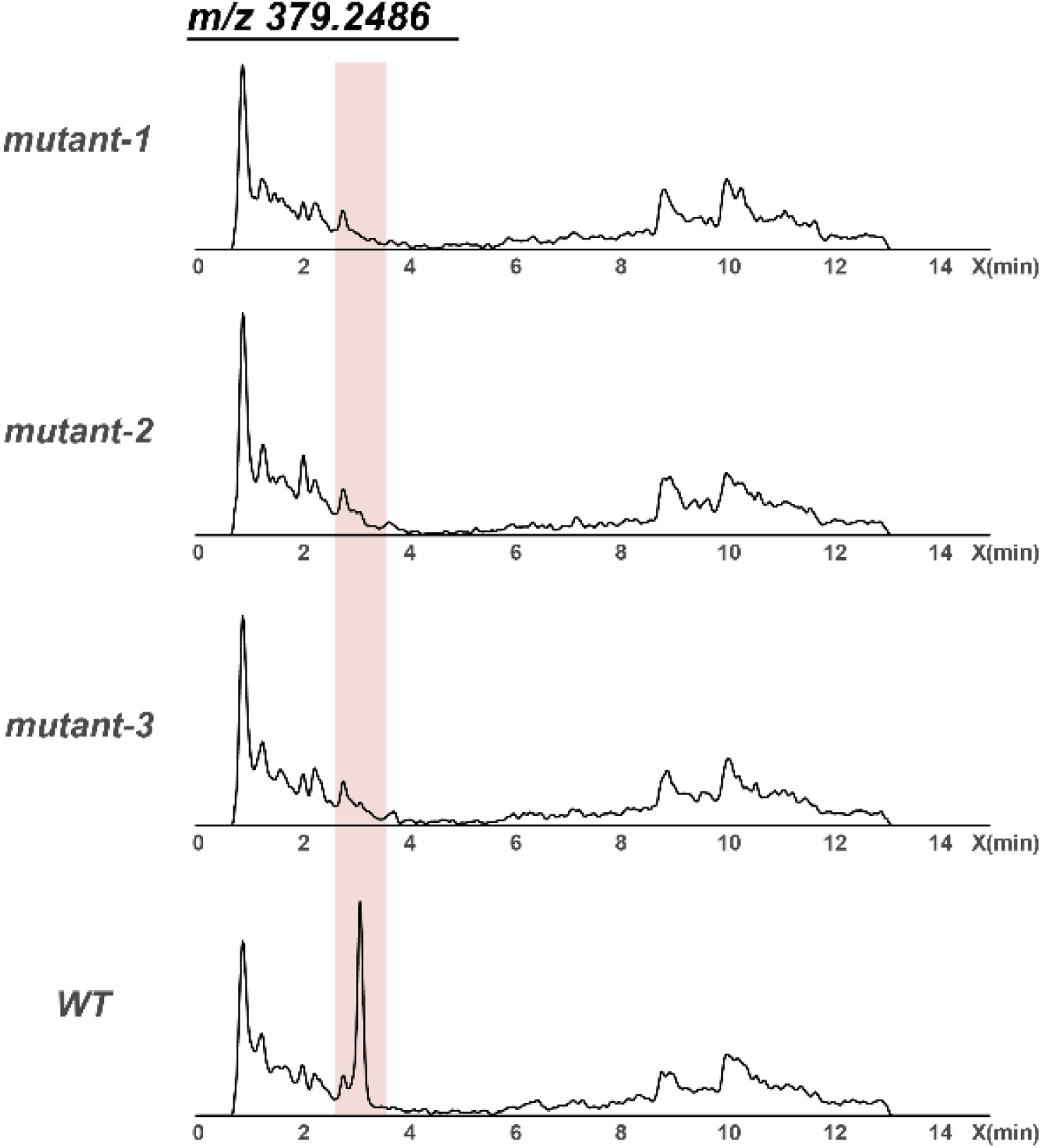
LC-MS analysis of metabolites extracted from CYP20 cultures of wild-type (WT) and mutant A. oryzae strains lacking the novel core gene. Extracted ion chromatograms (EICs) at m/z 379.2486 reveal WT-specific peaks (highlighted in red), absent in the mutant, indicating products of the novel gene.

**Supplementary Figure 10.**
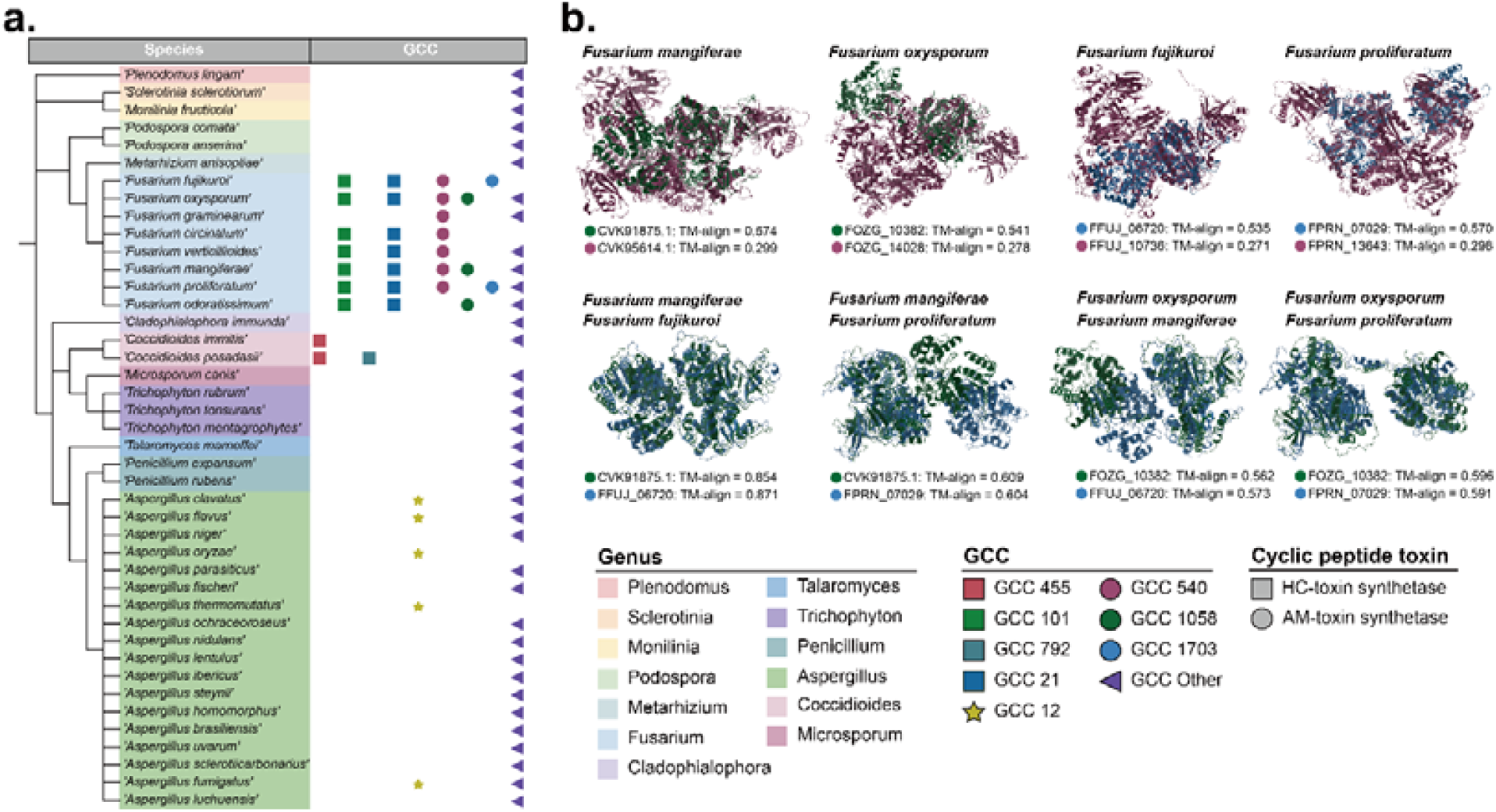
Landscape of phytotoxin candidate and potential BGCs. **a.** A phylogenomic tree built at the order taxonomic rank, colored by genus, with the corresponding GCC distribution shown in the right column. AMT GCCs (GCC540, GCC1058, GCC1703) are unique to *Fusarium*, HCT GCC is present in *Fusarium* and *Coccidioides*, GCC101 and GCC21 are exclusive to *Fusarium*, while GCC455 and GCC792 are specific to *Coccidioides*. The siderophore family GCC12 is found only in *Aspergillus*. **b.** Structural similarity of AM-toxin core genes in GCC540, GCC1058 and GCC1703 from *Fusarium*. Among the three AMT GCCs, GCC1058 and GCC1703 exhibit high core gene structural similarity, whereas their similarity to GCC540 is relatively low.

**Supplementary Figure 11.**
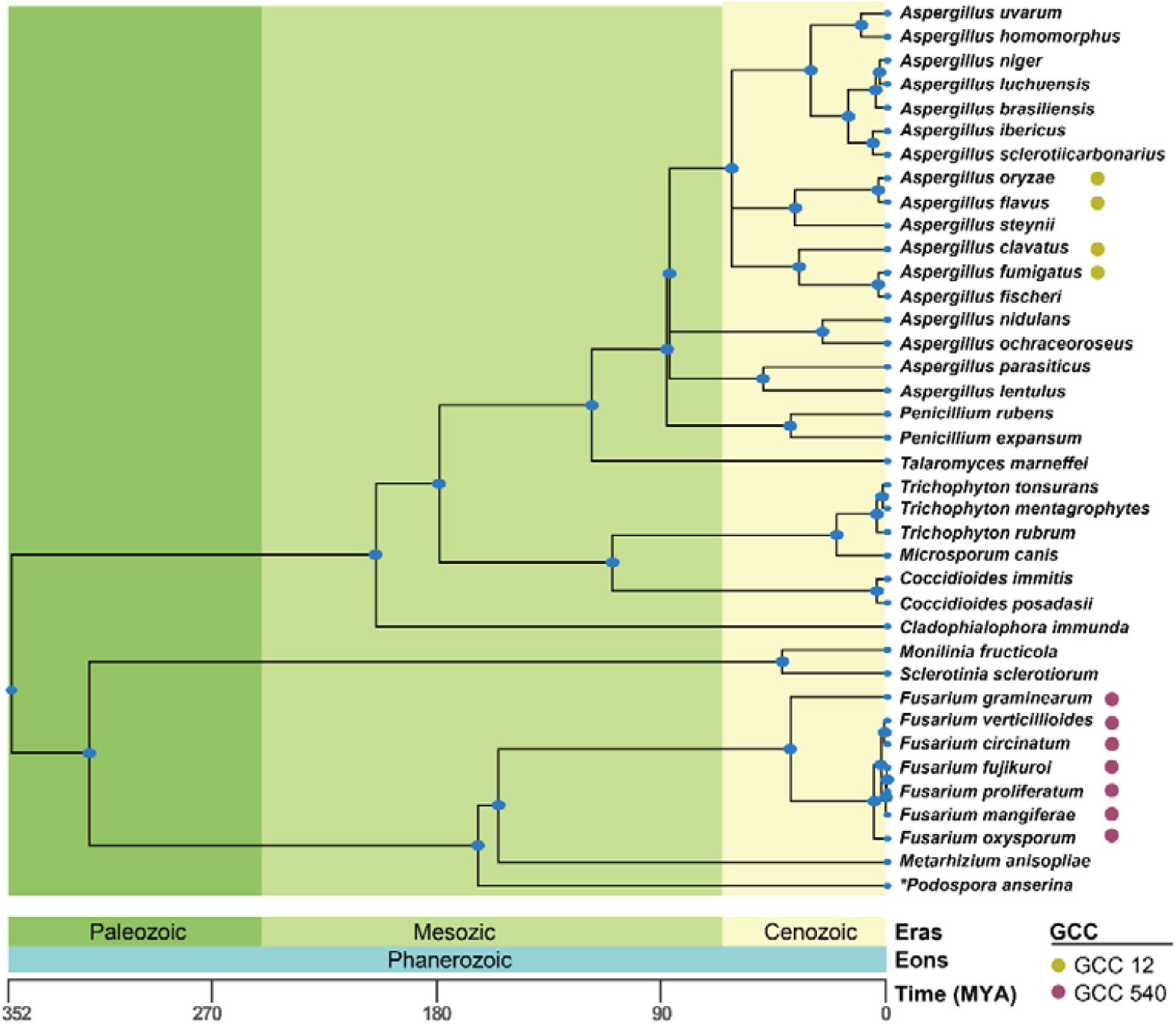
Evolutionary timeline of fungi. *Fusarium* and *Aspergillus* diverged approximately 352 million years ago, and during their long evolutionary history, each developed distinct GCCs: GCC540 in *Fusarium*, which is involved in AMT synthesis, and GCC12 in *Aspergillus*, which participates in siderophore synthesis. Despite their prolonged evolution and divergence into different functional roles, the core genes of GCC540 and GCC12 exhibit high structural similarity. This suggests that the emergence of these core genes predates the divergence of these two genera, with other species likely losing this core gene during their evolutionary process.

**Supplementary Figure 12.**
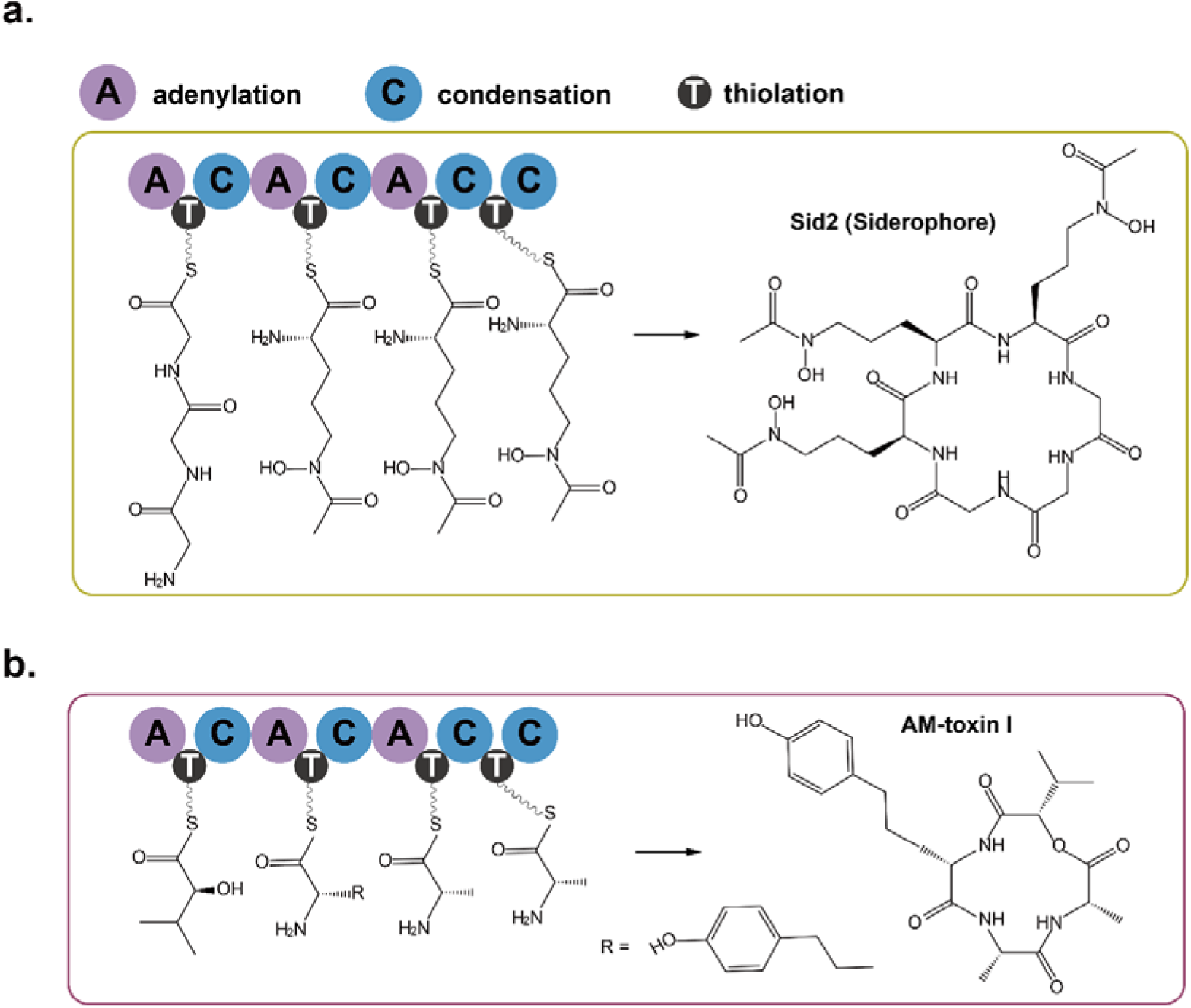
**The biosynthetic pathway of AM-toxin by GCC 540 and siderophore by GCC 12.**

